# Iterative transcription factor screening enables rapid generation of microglia-like cells from human iPSC

**DOI:** 10.1101/2022.06.03.494617

**Authors:** Songlei Liu, Li Li, Fan Zhang, Björn van Sambeek, Evan Appleton, Alex H. M. Ng, Parastoo Khoshakhlagh, Yuting Chen, Mariana Garcia-Corral, Chun-Ting Wu, Jeremy Y. Huang, Yuqi Tan, George Chao, John Aach, Jenny Tam, Elaine T. Lim, Soumya Raychaudhuri, George M. Church

**Affiliations:** Department of Genetics, Blavatnik Institute, Harvard Medical School, Boston, MA, USA; Wyss Institute for Biologically Inspired Engineering, Harvard University, Boston, MA, USA; Center for Data Sciences, Brigham and Women’s Hospital, Boston, MA, USA; Division of Genetics, Department of Medicine, Brigham and Women’s Hospital, Boston, MA, USA; Department of Biomedical Informatics, Harvard Medical School, Boston, MA, USA; Broad Institute of MIT and Harvard, Cambridge, MA, USA; Division of Rheumatology, Inflammation, and Immunity, Brigham and Women’s Hospital and Harvard Medical School, Boston, MA, USA; Radboud University, Nijmegen, the Netherlands; GC Therapeutics, Inc, Cambridge, MA, USA; Institute for Cell Engineering, Johns Hopkins University School of Medicine, Baltimore, MD, USA; Department of Molecular Biology and Genetics, Johns Hopkins University School of Medicine, Baltimore, MD, USA; Program in Bioinformatics and Integrative Biology and Departments of Neurology and Molecular, Cell and Cancer Biology, University of Massachusetts Chan Medical School, Worcester, MA, USA; NeuroNexus Institute, University of Massachusetts Chan Medical School, Worcester, MA, USA; Centre for Genetics and Genomics Versus Arthritis, Centre for Musculoskeletal Research, The University of Manchester, Manchester, UK

## Abstract

The ability to differentiate stem cells into human cell types is essential to define basic mechanisms and therapeutics, especially for cell types not routinely accessible by biopsies. But while engineered expression of transcription factors (TFs) identified through TF screens has been found to rapidly and efficiently produce some cell types, generation of other cell types that require complex combinations of TFs has been elusive. Here we develop an iterative, pooled single-cell TF screening method that improves the identification of effective TF combinations using the generation of human microglia-like cells as a testbed: Two iterations identified a combination of SPI1, CEBPA, FLI1, MEF2C, CEBPB, and IRF8 as sufficient to differentiate human iPSC into microglia-like cells in 4 days. Characterization of TF-induced microglia demonstrated molecular and functional similarity to primary microglia. We explore the use of single-cell atlas reference datasets to confirm identified TFs and how combining single-cell TF perturbation and gene expression data can enable the construction of causal gene regulatory networks. We describe what will be needed to fashion these methods into a generalized integrated pipeline, further ideas for enhancement, and possible applications.

## Introduction

Recent advances and applications of single-cell assays, exemplified by collaborative efforts such as the Human Cell Atlas (HCA)^1^, have begun to provide a comprehensive view of cell types and cellular states within the human body. Such maps are crucial for understanding human development and diseases. From a synthetic biology perspective, these maps can be mined for promising targets for cell fate engineering, with significant implications for disease modeling, cell therapy, and regenerative medicine. Previously, we reported the construction of a comprehensive human transcription factor library (TFome)^2^. In that study we used an unbiased approach for screening differentiation and identified 290 transcription factors (TFs) that induced differentiation of human induced pluripotent stem cells (hiPSCs) into various cell types. While the unbiased screening method led to many interesting discoveries, it does not guarantee the generation of any particular cell type. For those wishing to differentiate stem cells into a specific cell type of interest for studying diseases and creating therapeutics, the availability of experimental and computational pipelines for the identification of TFs to produce target cell types would be of great benefit. In this study, we picked a target cell type for which TF-based differentiation method has not yet been found, the microglia, for a proof-of-principal for developing new screening methodologies.

Microglia are the resident immune cells of the brain, which originated from erythro-myeloid progenitors (EMPs) in the yolk sac^3,4^. They play important and diverse roles in brain development and maintaining homeostasis^5–9^. Recent studies have demonstrated the link between neuroinflammation and neurodegenerative disease, such as Alzheimer’s Disease (AD)^10,11^, and along these lines, microglia have been shown to be an important cell type in AD and other neurodegenerative diseases^12–15^. However, functional studies to define therapeutics targeting human microglia have been greatly hindered by the limited availability of human brain biopsies^16,17^. The supply issue cannot simply be mitigated by using murine models, because differences between human and mouse microglia limit the transferability of knowledge^18,19^. Producing human microglia-like cells from hiPSCs might fill this gap. Several studies have accomplished this goal through a process of embryoid body formation, growth factor treatment, and, in some cases, co-culturing with neurons^20–28^. These protocols draw inspiration from the natural developmental stages of microglia and have timelines ranging from 30-74 days. As the effects of extrinsic factors on cell fate are frequently mediated by TFs, and building on top of our group’s and others’ prior research in using TFs to accelerate differentiation^2,29–31^, we hypothesized that direct manipulations of TF expression could differentiate hiPSCs to microglia in a shorter timeframe.

In this study, we conducted two sequential iterations of pooled TF screening. Each round of screening involves creating a barcoded TF library, pooled transfection into iPSCs for inducing differentiation, and subsequent single-cell transcriptome analysis. From the analysis, TFs are ranked by their ability to induce microglial gene expression, and the top hits then characterized for their ability to induce differentiation into microglia (**Figure 1a, 2a**). We identified a TF combination that produced cells transcriptionally resembled microglial cells in four days without the need for media exchange. We demonstrated that these TF-induced microglia-like cells (TFiMGLs) shared molecular and functional features of human primary microglia. Our barcoding and amplification strategy allow for simultaneous detection of cell and TF barcodes from the high throughput single-cell experiments, thus empowering us to analyze the regulatory relationships between TFs and other genes. We also constructed a human single-cell transcriptome reference by integrating publicly available scRNA-seq datasets of 225 samples representing 59 tissue types, to which TF differentiated single cells could be mapped. We expect the methodology described in this study to be broadly adopted to cell fate engineering and enable researchers to more effectively select TFs to generate novel iPSC-derived cell types.

**Figure 1.**
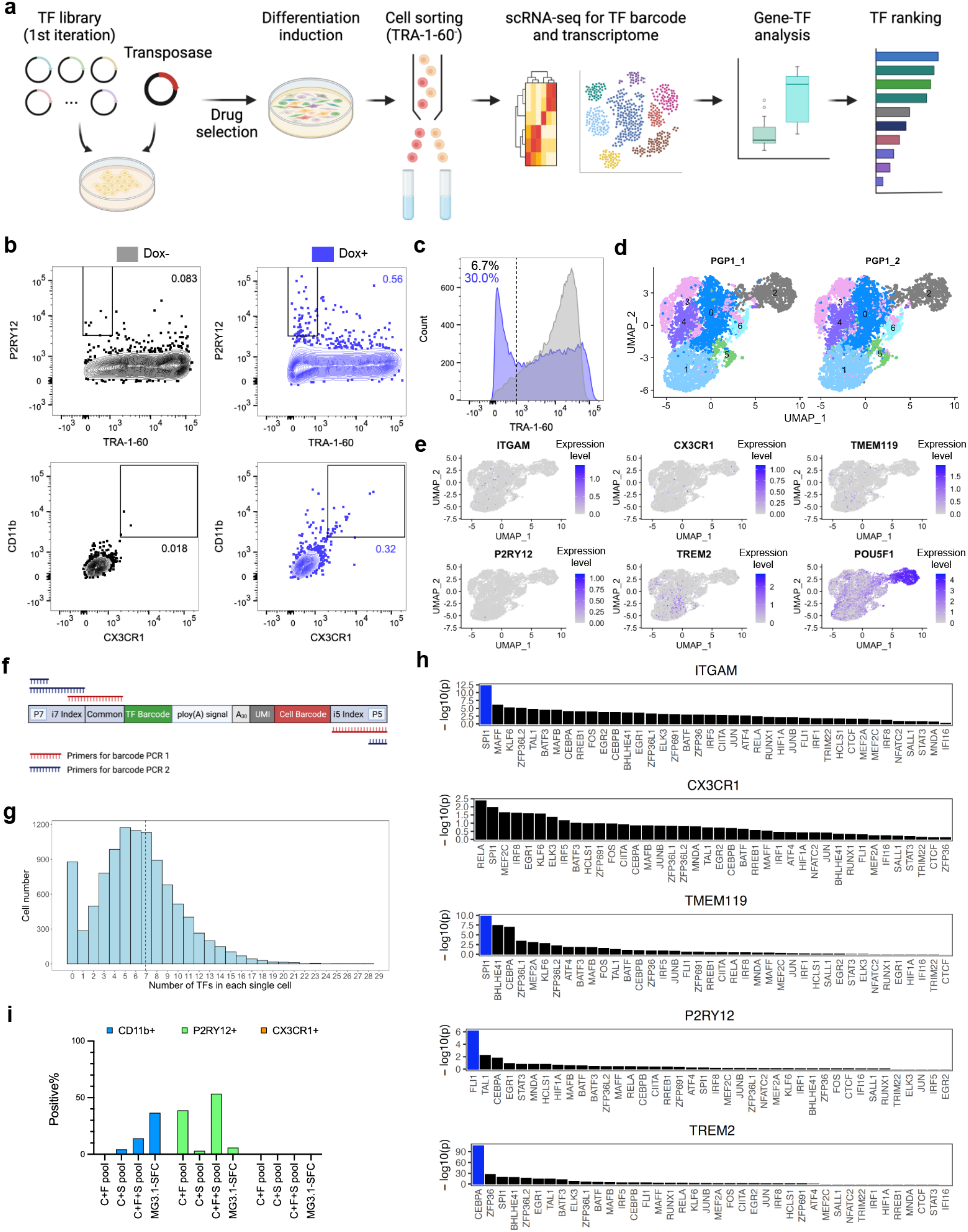
First round of pooled screening identified initial TFs for inducing microglia gene expression. (**a**) Workflow of the first pooled TF screen. (**b**) Flow cytometry analysis of stem cell (TRA-1-60) and microglia (P2RY12, CD11b, CX3CR1) proteins in the PGP1 + 40 TF pool before and after Dox induction. (**c**) Cells with low TRA-1-60 expression in the Dox+ group were sorted for scRNA-seq. (**d**) Clustering of two independently transfected and differentiated PGP1 iPSC pools. Colors represent clusters identified by Seurat at 0.3 resolution. (**e**) Expression of microglia (*ITGAM, CX3CR1, TMEM119, P2RY12, TREM2*) and spiked-in stem cell (*POU5F1*) gene in scRNA-seq. (**f**) Primer designs for co-amplification of TF and cell barcodes in 10x Genomics 3’ workflow. (**g**) Number of TFs per cell counted from normalized and binarized TF expression matrix. (**h**) Ranking of the 40 TFs after Wilcoxon rank sum test with the two tested groups being with or without microglia gene expression. Blue highlights top-ranking TFs. (**i**) Flow cytometry validation of top-ranking TFs for inducing microglia protein expression. C = CEBPA, F = FLI1, S = SPI1. “Pool” means pooled transfection, no polycistronic cassette used.

**Figure 2.**
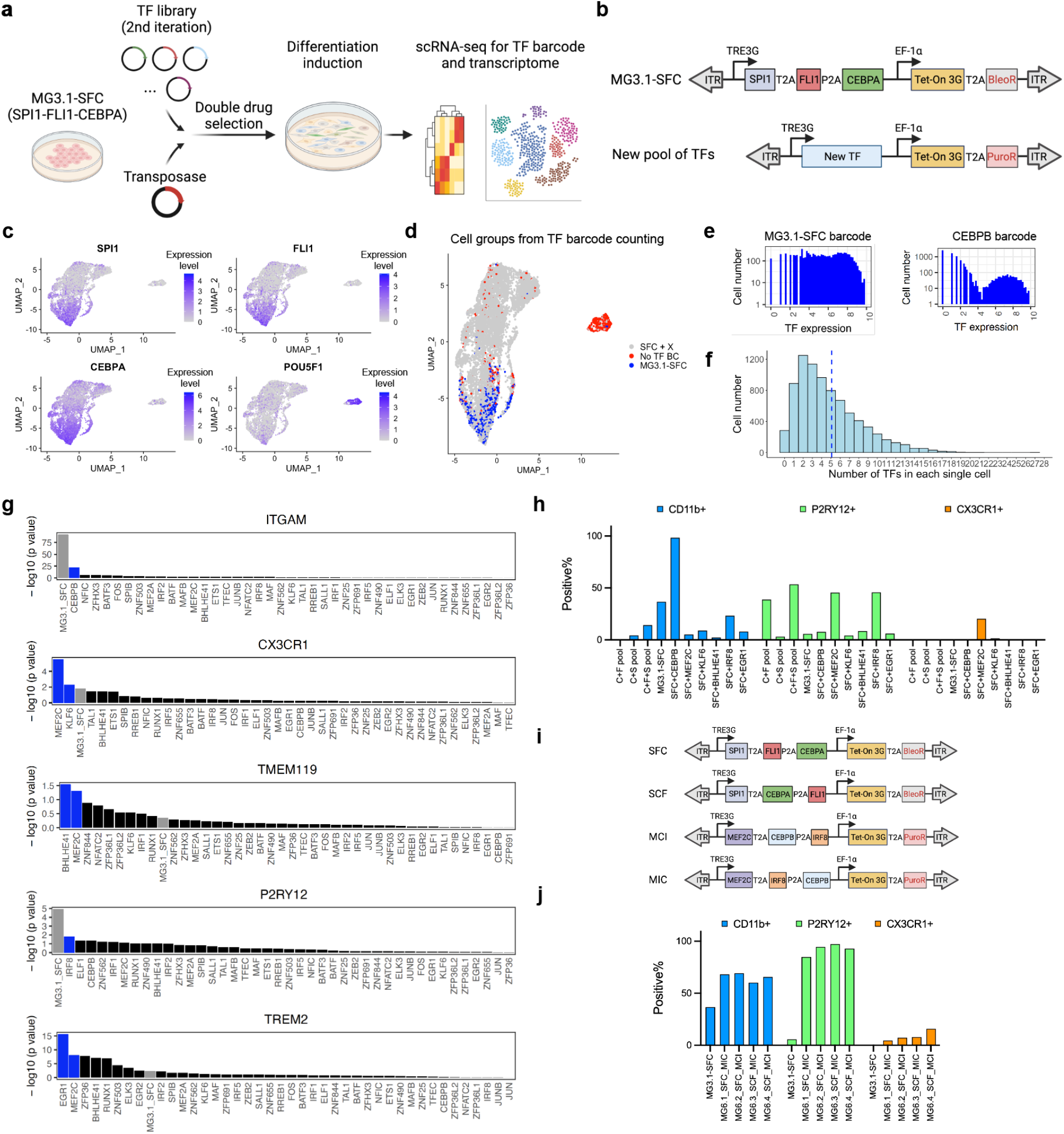
Second iteration of pooled TF screen using MG3.1-SFC as baseline identified additional TFs for improved microglia differentiation. (**a**) Workflow of the second pooled TF screen. (**b**) Polycistronic cassette design for performing dual-drug selection to achieve 3+X TF screen. (**c**) Normalized mRNA expression from the polycistronic cassette (*SPI1, FLI1, CEBPA*) and stem cells (*POU5F1*). (**d**) TF barcode counting enabled identification of stem cells (“No TF BC”), MG3.1-SFC and cells with additional TFs (“SFC+X”). (**e**) Example histograms of TF barcode raw counts in single cells. (**f**) Number of TFs per cell counted from normalized and binarized TF expression matrix. (**g**) Ranking of the 42 TFs after Wilcoxon rank sum test with the two tested groups being with or without microglia gene expression. Blue highlights top-ranking TFs. Grey highlights the SFC polycistronic cassette. (**h**) Flow cytometry validation of top-ranking TFs for improving microglia protein expression. (**i**) Polycistronic cassettes design for varying TF orders. (**j**) Flow cytometry analysis of different arrangements of the six-TF recipe in comparison with MG3.1-SFC.

## Results

### First round of pooled screening identified initial TFs for inducing microglia gene expression

To identify TFs that differentiate hiPSCs to microglia, our strategy is to first transfect and stochastically integrate a pool of TFs into the cells, followed by differentiation induction and single-cell RNA sequencing (scRNA-seq). From the scRNA-seq data, we use gene-expression profiles to determine which cells are differentiating into microglia, and identify the TFs that were transfected into these cells. To begin with, we surveyed previous literature on microglial development^3,6,8,32,33^, epigenetic and transcriptomic patterns^34–38^, and gene regulatory networks^39^. We shortlisted 40 TFs for the first pooled TF screening (**Supplementary Table S1**). We cloned each TF into the pBAN^2^ vector for genomic integration with PiggyBac transposase and doxycycline (Dox)-inducible expression. To distinguish between exogenous and endogenous TF transcripts, we added a 20-nucleotide (nt) barcode between the stop codon and the poly-A sequence of each TF (**Supplementary Figure S1**). We transfected the 40 TF vectors into 600,000 hiPSCs from a healthy donor (PGP1) with mass ratio between TF and transposase DNA being 4:1, with the assumption that each cell can uptake and integrate multiple TFs into their genome. After puromycin selection for TF-integrated cells, we induced differentiation by Dox for four days (**Figure 1a**). With flow cytometry we observed that 0.3-0.5% of the cells expressed microglial surface proteins, including CX3CR1, P2RY12, and CD11b (**Figure 1b**).

After four days of differentiation, we observed that 30% of the cells lost expression of a stem cell marker, TRA-1-60 (**Figure 1c**). To pinpoint which of the 40 TF(s) were inducing microglial gene expression, we sorted all differentiated (TRA-1-60 negative) cells for scRNA-seq (**Figure 1a**). We performed two independent transfections (**Supplementary Figure S2**), and spiked in 10% non-induced hiPSCs into each replicate as undifferentiated control during scRNA-seq. After observing reproducible differentiation between the two replicates (**Figure 1d**), we pooled the data together for downstream analysis. We observed expression of microglia genes (*ITGAM, P2RY12, CX3CR1, TMEM119, TREM2*), as well as a cluster of cells with high expression of *POU5F1*, marking stem cells (**Figure 1d, e**). Through amplicon sequencing of co-amplified TF and cell barcodes from cDNAs (**Figure 1f**), we quantified expression of exogeneous TF(s) in single cells in parallel (**Supplementary Figure S3**). In the Dox-induced cells, an average of 6.9 TFs (median 6, first quantile 4, third quantile 9) were expressed per cell. And 877 (8.5%) out of 10285 single cells had no TF expression, consistent with the 10% stem cell spike-in (**Figure 1g**).

We compared TF expression levels in cells with or without microglial RNA expression to identify TF(s) correlated with a higher expression of microglial genes. Presumably these TFs were the key TFs that had the potential to drive microglial differentiation. We identified three TFs likely to cause microglial gene expression: *SPI1, FLI1*, and *CEBPA* (**Figure 1h**). *SPI1*, which encodes PU.1 protein, is a known TF required for microglia development^32,33^. *CEBPA* is a known critical regulator for myeloid differentiation^40^. *FLI1*, while hasn’t yet been reported for microglial differentiation, has been reported to interact with *RUNX1*^41^ and *SPI1*^42^, where both TFs are indispensable for tissue-resident macrophage development^43^.

We wanted to understand if these TFs could lead to microglial differentiation individually, or if they needed to be used combinatorically. Individually expression of *CEBPA* and *FLI1* in hiPSCs led to almost complete cell death, indicating the need for additional TFs to stabilize the induced gene expression network. Expression of SPI1 alone was also ineffective, leading to only induction of CD11b in 3% cells (**Supplementary Figure S4**), indicating that multiple TFs are needed for the microglia differentiation.

We then tested combinations of TFs. Pooled transfection of *CEBPA+FLI1* (“C+F pool”) or *CEBPA+SPI1* (“C+S pool”) led to improved microglial marker expression, while *CEBPA+FLI1+SPI1* (“C+F+S pool”) produced the most positive cells, reaching 14% CD11b+, 54% P2RY12+ after four days (**Figure 1i**). However, we observed no expression of CX3CR1, a chemokine receptor important for microglia activation and migration^44,45^. Because pooled transfection and PiggyBac integration of three plasmids does not guarantee that every cell expressed all three TFs, we built polycistronic expression cassettes by linking the TFs with 2A peptides. A previous study reported that the gene position in the cassette affects their relative expression level, with the first gene being the highest-expressed^46^. Therefore, we arranged the TFs in different orders (**Supplementary Figure S5**). We named the construct by ordering letters corresponding to each TF corresponding to their order on the plasmid. For example, ***S***PI1-T2A-***F***LI1-P2A-***C***EBPA was named “MG3.1-SFC”. Transfection and induction of the MG3.1-CFS and FCS cassettes both led to dramatic cell death by day 4, consistent with the previous observation that sole CEBPA and FLI1 expression caused cell death. MG3.1-SFC, which positioned *SPI1* at the front, produced cells expressing two microglial genes, CD11b+ and P2RY12+ cells (37% and 6% of cells, respectively), but no *CX3CR1* expression (**Figure 1i**). The differences between cells derived using the MG3.1-SFC cassette and the microglia-like cells from the C+F+S pool are potentially due to different dosages of the TFs. While individual cells within the C+F+S pool may have expressed variable dosage combination of the three TFs, MG3.1-SFC likely induced a fixed dosage ratio for all cells. Critically, the lack of CX3CR1 expression and the low percentage of CD11b- and P2RY12-positive cells from all 3-TF conditions indicated that additional TFs were needed for efficient microglia differentiation from hiPSCs.

### Second iteration of pooled TF screen using MG3.1-SFC as baseline identified additional TFs for improved microglia differentiation

Recognizing that the three TFs identified in the first iteration were in themselves inadequate to differentiate microglia, we pursued a second iteration of our screen. To build upon the hits from the first pooled screen and identify additional TFs essential for microglia differentiation, we performed a second pooled screen. For this iteration, we used the top three TFs from the first iteration as baseline and tested the addition of other TFs (3+X) (**Figure 2a**). To determine what TFs should be included in the second pool, we performed a bulk RNA-seq analysis for MG3.1-SFC and compared it with published data on human primary microglia (GSE89189, GSE99074)^21,35^. Based on differential gene expression analysis using DESeq2^47^, we first picked 25 TFs included in the first pool that still had lower expression levels in MG3.1-SFC than primary microglia. We then included six new TFs that are significantly higher in primary microglia. By looking at which TFs regulate the genes that have lower expression in MG3.1-SFC using Molecular Signatures Database (MSigDB)^48^ regulatory target gene sets, we included *IRF2* and *ELF1*. Additionally, using CellNet^49^, a computational tool that can classify bulk transcriptomic data and predict missing gene regulators, we added six more TFs to the pool. Lastly, referring to a recent single-cell study on fetal microglia development^50^, we added *SPIB, ETS1* and *ELK3*. Thus, the second TF pool contained a total of 42 TFs (**Supplementary Table S2**).

To ensure each cell expressed both the *SPI1-T2A-FLI1-P2A-CEBPA* cassette and additional TFs from the second pool, we cloned the SFC cassette into a bleomycin-resistant vector and the new TF pool into a puromycin-resistant vector (**Figure 2b**). We transfected 600,000 PGP1 hiPSCs in duplicates and performed dual-drug selection for genomic integration. After four days of TF expression, we performed the same process of scRNA-seq and TF barcode amplicon sequencing as in the first iteration (**Supplementary Figure S6**). As controls, we spiked in 5% undifferentiated hiPSCs and 10% MG3.1-SFC during single-cell encapsulation to mark the differentiation starting point of two iterations. We applied this approach to two pools of cells.

When we analyzed TF barcodes in this experiment, we observed two clusters of cells on UMAP that corresponded to hiPSCs and MG3.1-SFC, while also showed new clusters of cells that express additional TFs (**Figure 2c, d**). When we counted TF barcodes, we observed that out of the total 8051 single cells from two independent transfections, 284 (3.5%) cells had no TF barcode and 613 (7.6%) cells had only the barcode for MG3.1-SFC. On average each cell expressed five TFs (median 4, first quantile 3, third quantile 7) (**Figure 2e, f, Supplementary Figure S7**), with most cells (88.9%) expressing the SFC cassette plus at least one other TF.

To determine which of the new TFs lead to improved microglia differentiation, we analyzed their effects on microglial gene expression. We were especially interested in increasing expression of *CX3CR1*, which was not expressed in MG3.1-SFC. We observed significantly higher (p < 0.01) number of *MEF2C* and *KLF6* barcode in cells expressing *CX3CR1* (**Figure 2g**), suggesting their ability to induce *CX3CR1* expression. It’s worth noting that *MEF2C* was also present in the first screening but failed to reach significance for upregulating *CX3CR1*, indicating the use of the SFC cassette as baseline enabled other influential TFs to be found. *MEF2C* also reached high ranking for *TMEM119* (**Figure 2g**). The SFC cassette ranked at the top for inducing *ITGAM* and *P2RY12* expression, which is expected from the results of the first iteration. In this round of screening, *CEBPB* and *IRF8* emerged as high-potential TFs for promoting *ITGAM* or *P2RY12* expression (**Figure 2g**). From this second pooled TF screening, additional TFs of interest found were *MEF2C, CEBPB, IRF8, KLF6*, and *BHLHE41*.

To validate that these additional TFs can promote microglial gene expression, we individually expressed them in addition to SFC (SFC+1). When compared with MG3.1-SFC, SFC+CEBPB increased the percentage of CD11b+ cells from 37% to 98% (**Figure 2h**) but led to more cell death at day 4. SFC+MEF2C and SFC+IRF8 increased P2RY12 expression from 6% to 45% (**Figure 2h**). Most importantly, SFC+MEF2C and SFC+KLF6 increased CX3CR1+ cells from 0% to 20% and 2% respectively (**Figure 2h**). These results agreed well with the predictions from single-cell TF barcode analysis, indicating the validity of using pooled TF screening for inferring causality between TF and target gene expression.

To see if microglia differentiation can be further promoted by delivering more TFs to each cell, we chose the three TFs from the SFC+1 experiment that led to highest increase in percentage of microglial gene-expressing cells, CEBPB, IRF8, and MEF2C, to add to the SFC set. We combined MEF2C, CEBPB and IRF8 into polycistronic cassettes. Because MEF2C demonstrated ability to induce both CX3CR1 and P2RY12, we put it in the first place and varied the position of CEBPB and IRF8, producing two cassettes: MIC and MCI (**Figure 2i**). We also varied the position of FLI1 and CEBPA in the first construct to produce SFC and SCF, keeping SPI1 in the front to avoid excessive cell death during differentiation. We tested all four combinations of the two 3-TF cassettes (SFC-MIC, SFC-MCI, SCF-MIC, SCF-MCI) for their ability to induce microglia differentiation (**Figure 2i**). Encouragingly, all 6-TF cocktails produced cell pools with increased expression of microglial proteins when compared with MG3.1-SFC (**Figure 2j**). We observed that the most effective combination was MG6.4-SCF-MCI, resulting in 66% CD11b+, 93% P2RY12+ and 16% CX3CR1+ cells at day 4, compared with 37%, 6%, and 0% respectively for MG3.1-SFC, the baseline of the second iteration. These results highlighted the value of the second iteration and demonstrated the utility of iterative TF screening for cell fate engineering. We term cells differentiated using MG6.4-SCF-MCI Transcription Factor-induced MicroGlial-Like cells, or TFiMGLs.

### TFiMGLs are phagocytic, responsive to disease-relevant stimulation, and share molecular signatures with primary microglia

To determine the differentiation dynamics of TFiMGLs, we performed bulk RNA-seq analysis of the cells on 0, 1, 2, 3, 4, 6 days post induction of the six TFs. We observed a rapid induction of all six TFs on day 1, and they reached plateau on day 2 (**Figure 3a**). This accompanied a quick downregulation of *POU5F1* on day 1 and followed by upregulation of microglial genes from day 2 and onwards (**Figure 3b**). Principal component analysis (PCA) reflected a similar trend, where a rapid differentiation occurred on day 1 and 2, followed by a gradual deceleration from day 3 to day 6 (**Figure 3c**). In a plot of PC1 and PC2, the day 4 and day 6 transcriptomes are close together, suggesting a stable window for functional investigation. For downstream characterizations of TFiMGLs, we chose to differentiate cells for 4 days.

**Figure 3.**
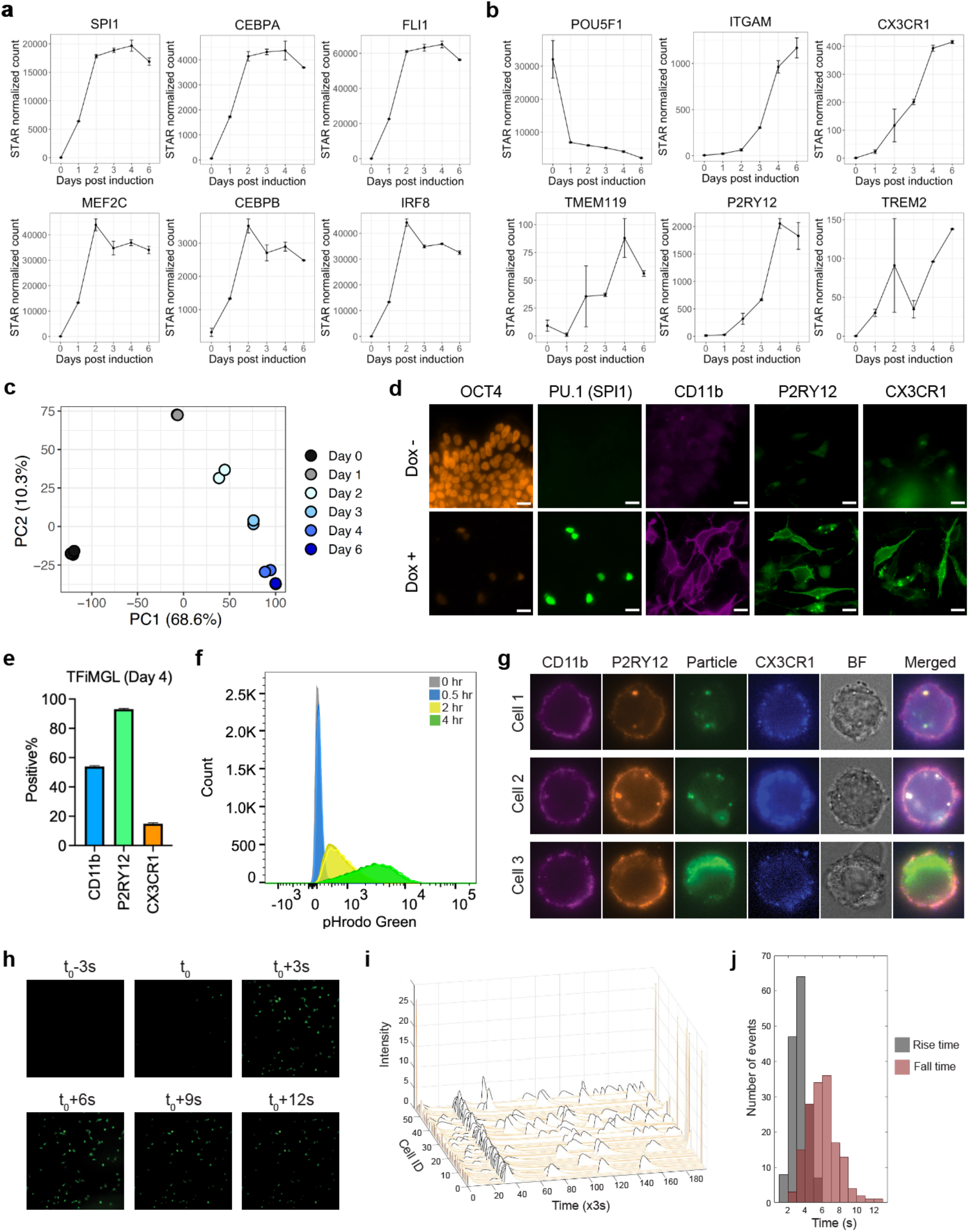
TFiMGLs differentiate quickly, are phagocytic and responsive to ADP stimulation. (**a**) Expression of the six induced TFs over time measured by bulk RNA-seq. n = 2 for each day. Error bar represents standard deviation. (**b**) Expression of stem cell (*POU5F1*) and microglia (*ITGAM, CX3CR1, TMEM119, P2RY12, TREM2*) genes over time measured by bulk RNA-seq. (**c**) PCA plot for the transcriptome of TFiMGLs (MG6.4) over time. (**d**) Immunofluorescence of stem cell (OCT4), Dox-induced (PU.1), and microglia (CD11b, P2RY12, CX3CR1) proteins on day 4. Scale bar: 20 μm. (**e**) Flow cytometry quantification of microglia protein expression on day 4 (n=3). (**f**) Flow cytometry analysis of the uptake of pHrodo-labeled S. aureus Bioparticles over time (n=3). (**g**) Microscopy analysis of particle uptake combined with microglia surface protein staining. (**h**) Calcium imaging with Fluo-4 after stimulation with 150 μM ADP and peak quantification. Images taken once every three seconds. ADP was added at t_0_. (**i**) Quantification of fluorescent signals from all cells in the field of view in panel h over a period of 10 minutes. (**h**) Peak dynamics analysis shows a fast rise and slow decay pattern of the intracellular calcium concentration.

Brightfield microscopy analysis of TFiMGLs revealed rapid morphological change from day 1 to day 6 (**Supplementary Figure S8**). Immunofluorescence analysis confirmed the loss of pluripotency marker OCT4 and expression of key microglial proteins: CD11b, P2RY12, and CX3CR1 (**Figure 3d**). TFiMGLs demonstrated reproducible differentiation between replicates, with 53.9 ± 0.57% (SD, n=3) CD11b+, 93.1 ± 0.50% (SD, n=3) P2RY12+ and 14.8 ± 0.68% (SD, n=3) CX3CR1+ cells (**Figure 3e**).

As brain resident macrophages, microglia play important roles in brain development and homeostasis. Microglia’s abilities to respond to signals related to degenerating neurons and phagocytose are integral parts of their function^21^. To investigate if TFiMGLs could mimic the phagocytosis function of microglia, we incubated TFiMGLs with pHrodo green labeled S. aureus particles for 0.5, 2 and 4 hours, and performed flow cytometry and microscopy. While 0.5-hour incubation showed minimal phagocytosis activity, nearly all cells were positive for pHrodo green at 2 hours and the intensity grew even stronger at 4 hours (**Figure 3f, Supplementary Video S1-2**). Microscopy analysis at 4 hours with co-staining of microglia surface proteins confirmed the intracellular position of these particles (**Figure 3g**). ADP is one of the substances released from injured neurons and works as an signal to stimulate microglial responses^51,52^. To study if TFiMGLs are responsive to ADP stimulation, we incubated TFiMGLs with calcium indicator Fluo-4 and then stimulated with ADP containing media. We imaged the cells at a three-second interval. We observed a rapid increase in calcium signal when ADP was added (**Figure 3h, i, j; Supplementary Video S3**), suggesting TFiMGLs are responsive to ADP stimulation.

To assess how accurately TFiMGLs recapitulate the transcriptome of human microglia, we compared TFiMGLs bulk-RNA-seq data to previously published bulk RNA-seq data for human primary microglia and iPSC-derived microglia (GSE89189, GSE99074)^21,34,35^. To minimize potential batch effects that might hinder meaningful comparison between datasets, we aligned all raw FASTQ files to the same reference genome and applied a negative binomial regression-based batch effect correction method, ComBat-seq^53^, before downstream analysis. From the PCA analysis, we observed that the transcriptomes of TFiMGLs from days 2-6 more closely resembled primary microglia of different sources than to iPSCs or hematopoietic progenitors (HPCs), suggesting a successful microglial fate induction (**Figure 4a**). We also observed that TFiMGLs were distinct from monocytes or dendritic cells, two related cell types from the myeloid lineage. To investigate if TFiMGLs express genes that are enriched in primary microglia, we performed Gene Set Enrichment Analysis (GSEA)^54^ on TFiMGLs versus iPSCs using two microglial gene collections from the MSigDB derived from human brain scRNA-seq (M40168; M39077). We observed significant positive microglial gene enrichment scores using both collections (M40168: score = 0.72, p-value = 9.01e-10, gene set size = 313; M39077: score = 0.66, p-value = 9.01e-10, gene set size = 391), indicating TFiMGLs upregulated those microglia-enriched genes (**Figure 4b**). To further investigate if TFiMGLs achieve transcriptomic similarity to primary microglia, we also used a previously published collection of 881 microglia-enriched genes^34^ to cluster the samples from **Figure 4a**. While the transcriptome of day-1 TFiMGLs clustered closer to iPSCs, day-2 and later TFiMGLs clustered closer to primary microglia, with key microglial genes increasingly upregulated by the day (**Figure 4c**). Similar to what we saw in **Figure 4a**, TFiMGLs were distinct from monocytes or dendritic cells by measuring the 881 genes. These analyses demonstrate that the TFiMGLs have lost iPSC-like identity and now closely resemble microglia.

**Figure 4.**
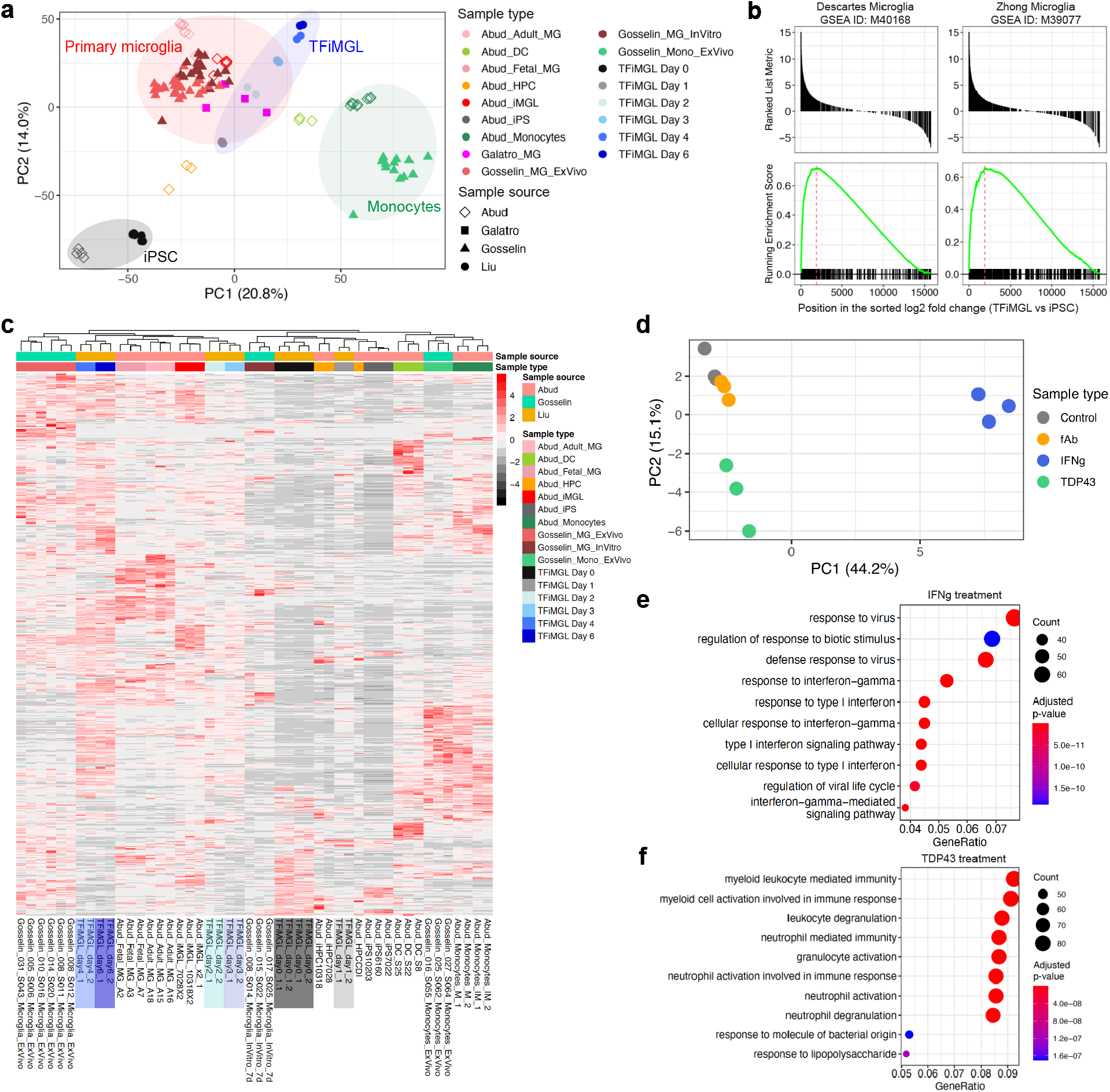
Transcriptome analysis of TFiMGLs on different days and under disease relevant stimulations. (**a**) PCA of bulk RNA-seq data from multiple sources containing primary microglia. MG: Microglia; DC: dendritic cell; HPC: hematopoietic progenitor; iMGL: growth factor-induced microglia-like cell; Mono: monocyte; iPS: induced pluripotent stem cell. (**b**) GSEA of TFiMGLs versus iPS using two microglia marker gene sets from MSigDB: M40168 and M39077. (**c**) Heatmap and clustering with 881 microglia specific genes previously reported (Ref. Gosselin). (**d**) PCA of TFiMGLs’ transcriptome after 24 hours treatment with IFNγ, fAβ, or TDP43. (**e**-**f**) Pathway analysis of significantly differentially expressed genes after treatment with IFNγ or TDP43.

Microglia are able to respond to signals indicating brain infection and inflammation, such) as IFNγ, beta amyloid, and TDP-43. IFNγ is a known activator of microglia secreted by TL lymphocyte^55^. Beta amyloid (Aβ) is a key molecule in AD pathology which has been shown to elicit microglia response^56^. TDP-43, whose aggregation is considered a hallmark of ALS and is present in the vast majority of amyotrophic lateral sclerosis (ALS) patient, had also been also I shown to activate microglia^57^. To investigate how TFiMGLs respond to IFNγ, fibrillar Aβ (fAβ) and TDP-43, we treated TFiMGLs in triplicates with each of the three molecules for 24 hours and harvested cells for RNA-seq. PCA analysis revealed transcriptomic changes in the IFNγ and TDP-43 treated group, while the fAβ-treated group showed minimal differences (**Figure 4d, Supplementary Figure S9**). We confirmed fAβ formation by conducting an in vitro amyloid fibrillation experiment that showed the Aβ peptide could form fibrils after 1 hour of incubation) (**Supplementary Figure S10**). Pathway analysis of differentially expressed genes from the IFNγ treated group included “response to virus” and “response to bacterium” (**Figure 4e, Supplementary Figure S11**), corresponding to the role of IFNγ production as a response to infection. Top upregulated genes by IFNγ included CXCL10, CXCL11, IRF1 and IL18BP (**Supplementary Figure S11**), which aligns with the IFNγ response genes revealed by an independent single-cell level human-derived macrophage stimulation study^58^. For the TDP-43 treated cells, top differentially regulated pathway included “myeloid leukocyte mediated immunity” and “myeloid cell activation involved in immune response” (**Figure 4f, Supplementary Figure S12**), demonstrating that TFiMGLs were activated by the TDP-43 treatment. Collectively, these results suggest that TFiMGLs exhibited microglia-like responses to infection- and ALS-related stimulations.

### Single-cell atlas reference mapping confirmed microglia-like fate induction

There is the opportunity to map cell states precisely leveraging advances in single-cell analysis technologies. Up until this point, we have been using a group of cellular markers to determine cell type. While this is a common practice in both primary human tissue and stem cell differentiation studies, it is now possible to define cell types based on more comprehensive molecular profiles, including the whole transcriptome. We aspired to develop a strategy that might be generally applicable to leveraging single-cell atlas data to guide cell fate engineering efforts.

To achieve this goal, there are several important prerequisites: 1) reference single-cell data sets from all developing human tissue types, capturing different developmental stages; 2) data integration methods to combine these different reference data sets to create a comprehensive cell atlas; 3) reference mapping methods to project iPSC-derived cell data onto the combined reference and quantitative assessment of similarity to differentiating cell classes; 4) existence of perturbation libraries that are scRNA-seq compatible, which include but not limited to open reading frame (ORF) and CRISPR libraries. While advances have been made and continue for #2-4, incomplete reference data continues to be an acute problem.

To explore this idea of atlas-guided cell fate engineering, despite limitations in available reference data, we re-examined the previously described two pooled TF screen for microglia differentiation. We built a refence data set by compiling scRNA-seq data from published datasets generated through 10x Chromium platform. In total, the final single-cell atlas contains 225 samples from 59 organ or tissue types, with a total of 1,004,650 single cells (**Supplementary Table S3**). The majority of the data was obtained from PanglaoDB^59^, where all raw reads from different studies were aligned and processed together. We added other data sets representing brain^60^ and endometrium^61^ which were under-represented in PanglaoDB. We carefully filtered all raw data downloaded from PanglaoDB for cell, gene, UMI number and mitochondria gene ratio (**Supplementary Table S4**). In its current form, the cells in the atlas are annotated according to their organ or tissue of origin; cellular level annotation for all 59 studies have yet to be defined.

To reduce batch variability across different studies, the data were integrated with Harmony^62^ (**Figure 5a**; **Supplementary Figure S13**). Qualitative assessment of the UMAP plots post-integration indicated co-clustering of cells in related tissues from different studies, exemplified by the overlap between “Primary brain” with “Embryo forebrain” datasets, as well as between the “Pancreatic islets” with “Pseudoislets” datasets (**Supplementary Figure S13**). To visualize where microglia are located on this map, we acquired cell annotation information for the “Primary brain” dataset^60^. We observed a cluster containing microglia on the right of the UMAP plot (**Figure 5b**). To see if any of the TF differentiated cells can be mapped closely to microglia, we projected the scRNA-seq data from the two pooled TF screens onto the integrated atlas using Symphony^63^ **(Figure 5c, d**). In the projections, we observed 4.3% and 26.5% cells from the first and second screen being projected to the microglia-containing cluster. This increase in percentage is like because the top hits from the first screen, SPI1, FLI1, and CEBPA were the baseline for the second screen. Because we have identified SPI1, CEBPA, FLI1, MEF2C, CEBPB, and IRF8 as the inducers for differentiating iPSCs to microglia-like cells, we wanted to see if their barcode expression lead to co-localization with microglia on the reference atlas. We saw CEBPA barcode had high expression in the microglia cluster, as well as a few others (**Figure 5e**), indicating its ability to activate a broad range of genes. SPI1 barcode, on the other hand, had a distinct enrichment in the microglia cluster (**Figure 5f**), suggesting a microglia specific gene induction. We observed the co-expression of SPI1, FLI1, and CEBPA by the SFC cassette led to a strong mapping to the microglia cluster (**Figure 5g**). We also visualized the expression of FLI1, MEF2C, CEBPB, and IRF8 individually (**Supplementary Figure S14**). While they did not show obvious enrichment in microglia cluster themselves, combined reads from the all six TFs still showed a strong induction towards microglia (**Figure 5h**). This single-cell atlas reference mapping analysis confirmed microglial fate induction by the six TFs with a comprehensive comparison with 1,004,650 single cells from 59 organ or tissue types.

**Figure 5.**
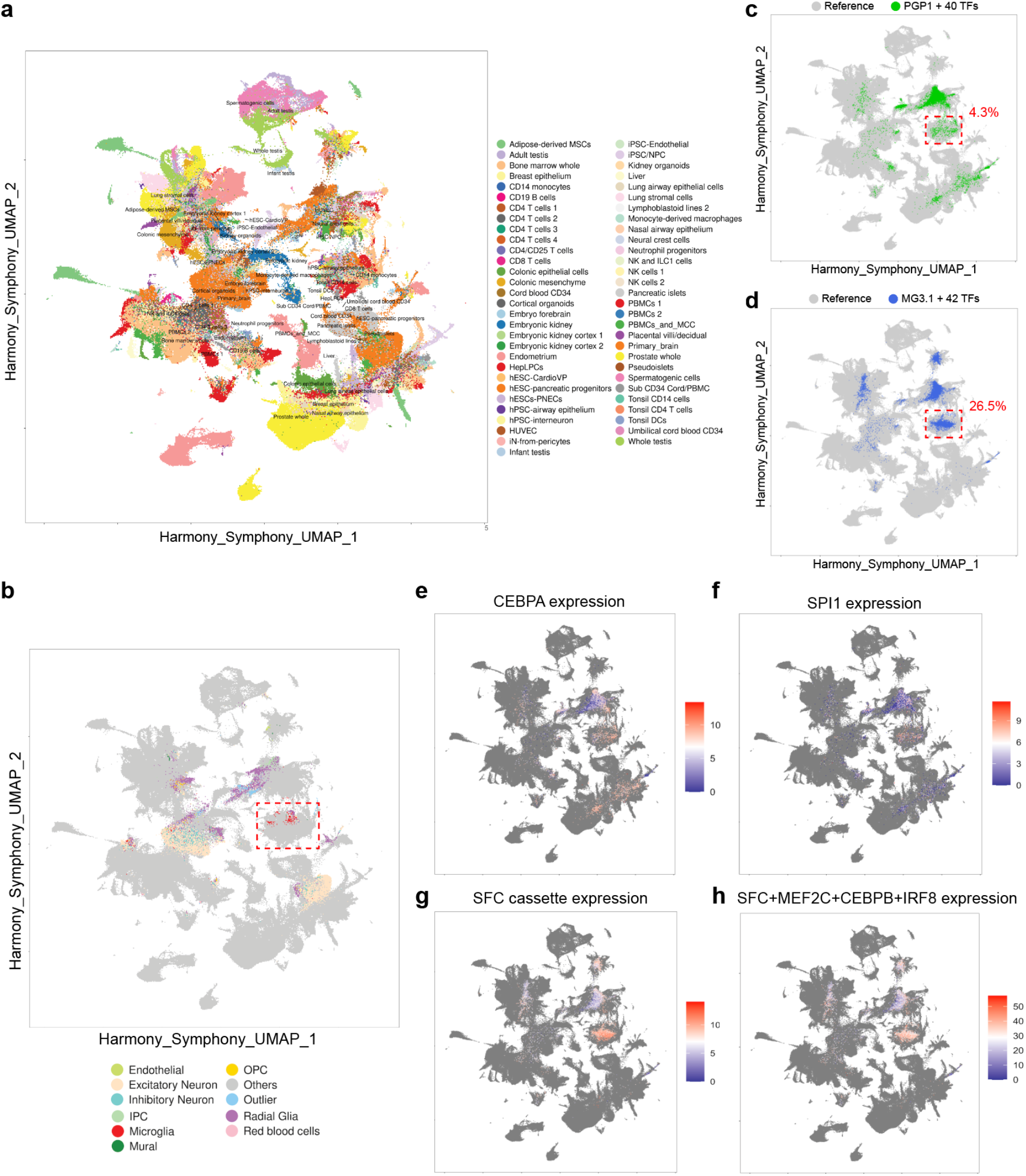
Single-cell atlas reference mapping confirmed microglia-like fate induction. (**a**) Harmony-integrated single-cell reference atlas with 225 samples from 59 organ or tissue types, with a total of 1,004,650 single cells. UMAP plot is colored by sample. Sample annotation is acquired by manual curation of each study. (**b**) UMAP is colored by the cellular level annotation from the “Primary_brain” dataset downloaded from “Organoid Report Card”. Red cells are microglia. The red dashed box highlighted the cluster containing microglia. (**c**) Symphony projection of cells from the first pooled screen (PGP1+40TFs) to the reference atlas. The red dashed box highlighted the 4.3% cells mapped to the microglia-containing cluster. (**d**) Symphony projection of cells from the second pooled screen (MG3.1+42TFs) to the reference atlas. The red dashed box highlighted the 26.5% cells mapped to the microglia-containing cluster. (**e**-**h**) Expression of key TF barcodes in the projected cells. CEBPA, SPI1, and the SFC cassette showed high expression in the microglia-containing cluster.

### Regression analysis revealed causal TF-gene regulatory relationships

Accumulating scRNA-seq data of human cell types provide ever-expanding information about what genes define a cell fate. As a result, a complete knowledge map of what TFs turn on what genes is instrumental for cell fate engineering. A tremendous amount of insight on this subject have been produced by computational methods for inferring TF-gene regulatory network (GRN)^64,65^ and databases based on TF-binding sites^66,67^, TF-gene co-expression^68,69^, and protein-protein interaction^70,71^. However, the only way to acquire a definitive causal TF-gene regulatory relationship map is by introducing the TF perturbations and observing their effects in a highly multiplexed way. Our two pooled TF screens combined with scRNA-seq readout enabled us to explore this exact idea. Taking advantage of our experimental design that captured both TF barcode and cellular RNA expression, and by implementing a stepwise regression model (Online Methods), we were able to construct GRNs from the two pooled screens (**Figure 6**). In our dataset, each TF transgene was represented by two distinct values: counts from barcode amplicon sequencing, and counts from their RNA molecules in scRNA-seq. Although the counts from scRNA-seq might contain reads from endogenous TF expression, the two measurements correlated well for most TFs (**Supplementary Figure S15**), the exception being a few TFs with low expression. We reasoned that TFs that had higher correlation between their barcode and RNA measurements demonstrated higher consistency between experiments, which were more reliable to produce accurate regression results using the two matrices. Thus, by selecting TFs had a correlation coefficient greater than 0.3 between the two measurements, we selected 18 TFs from the first iteration and 21 TFs from the second iteration for regression analysis. We observed extensive gene expression changes caused by CEBPA and the triple TF cassette MG3.1_SFC expression (**Figure 6a, f**), as well as slightly smaller networks from CIITA, SPI1, ERG2, JUN, CEBPB, ZFP36, and BHLHE41 (**Figure 6b-e, g, h, Supplementary Figure S16**). Among 672 edges, we observed 76% to be positive regulations. Some TFs (CIITA, SPI1, JUN) only showed positive edges in current thresholding conditions (Abs(coefficient) > 0.1 & -log_10_(p-value) > 20), indicating they were mostly activating other genes. Other TFs (CEBPB, ZFP36, BHLHE41) showed negative edges, indicating their repressive roles. We also observed several genes simultaneously connected with more than one TFs (**Supplementary Figure S17**). For example, RAB13, a membrane trafficking regulator, was upregulated by both CEBPA and CEBPB; HMGA1, a master regulator of chromatin structure, was downregulated by both BHLHE41 and CEBPA; FLNC, an actin crosslinking protein, was upregulated by JUN while downregulated by CEBPA. There are many more regulatory relationships we listed in detail from these two pooled screens (**Supplementary Table S5-6**). With larger perturbation libraries, higher-throughput scRNA-seq, and more scalable regression analysis methods, we believe a more complete knowledge map of causal TF-gene regulatory relationships could be built and greatly facilitate cell fate engineering efforts.

**Figure 6.**
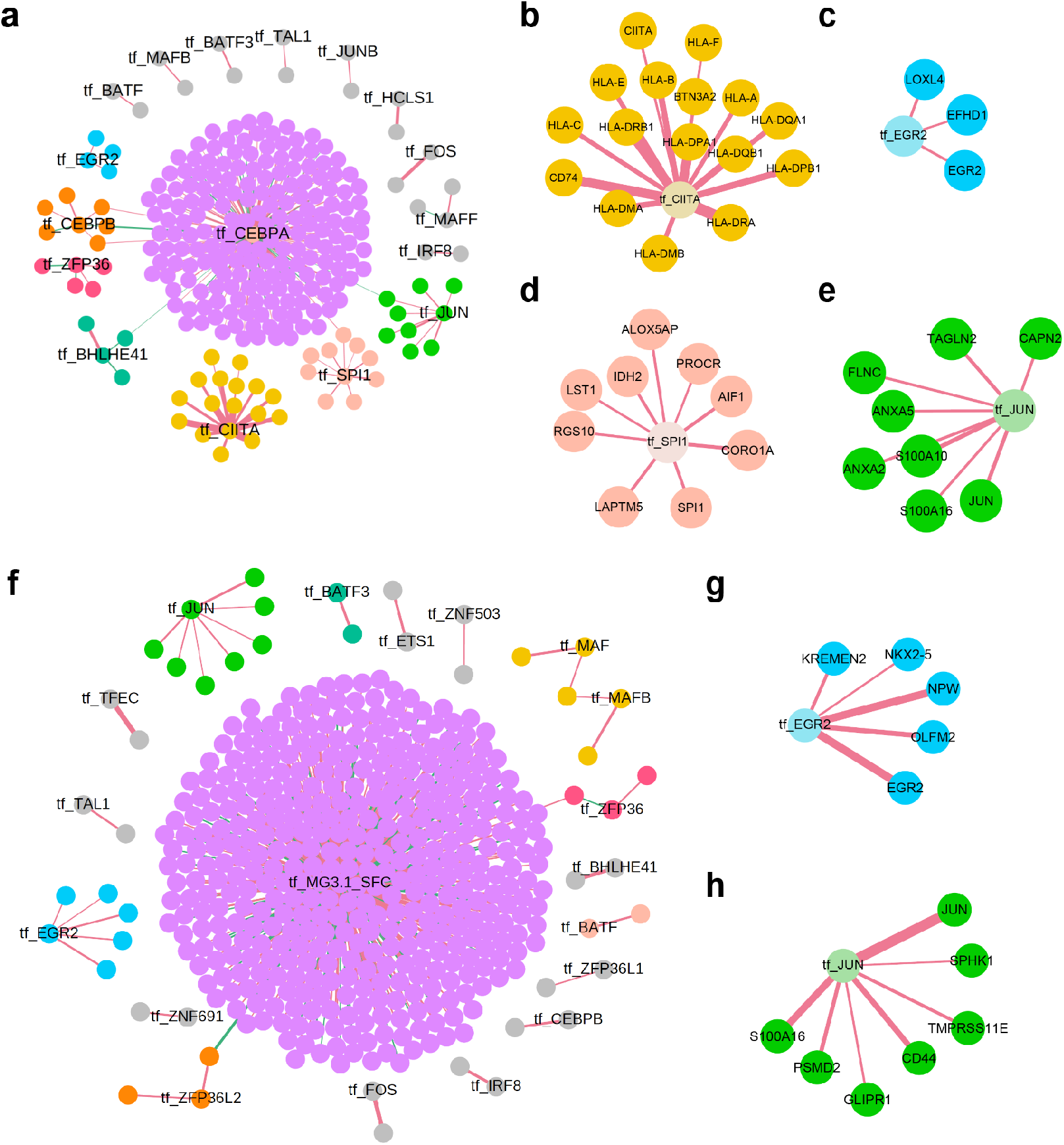
TF-gene regression analysis for studying causal gene-regulatory networks. Nodes’ label starts with “tf_” represent the TF captured by single-cell barcode sequencing. All other nodes represent genes captured by scRNA-seq. The width of edges is correlated with coefficient values. The larger the value the wider the edge. A red edge means upregulation while a green edge means downregulation. Edges were selected with these criteria: Abs(coefficient) > 0.1 & -log_10_(p-value) > 20. (**a**) Global network for the first pooled screen. Sub-network for (**b**) tf_CIITA, (**c**) tf_EGR2, (**d**) tf_SPI1, (**e**) tf_JUN. (**f**) Global network for the second pooled screen. Sub-network for (**g**) tf_EGR2 and (**h**) tf_JUN.

## Discussion

Differentiating human cell types from stem cells provides is essential for basic research and therapeutics development, especially when the desired cell types are not easily obtainable from accessible human tissues. Advances in the understanding of developmental biology has fueled the discovery and application of protocols to differentiate specific cell types from iPSCs. Some of this work has been translated into treatment strategies that are now being investigated with clinical trials for devastating diseases like age-related macular degeneration^72^ and type 1 diabetes^73^. Differentiated iPSCs have now also become routinely used in laboratories for studying disease mechanisms and testing drugs^74^. Recent global efforts on building single cell atlases of cellular development have expanded the knowledge of human development and diseases, but also present a key resource for cell fate engineering. Combined with technological advancements in genetic library construction, high-throughput screening and sequencing technologies, to the field is primed for investigating how to engineer cell fate in a systematic and multiplexed fashion, as performed in this work.

Our study demonstrated the feasibility of combining an iterative genetic library screen with high-throughput scRNA-seq for cell fate engineering. Using microglia, a cell type which previously did not have a TF-driven differentiation protocol, as a model target, we performed two iterations of our design-screen-validate workflow and identified SPI1, CEBPA, FLI1, MEF2C, CEBPB, and IRF8 as a potent recipe for driving microglia differentiation from hiPSCs within 4 days, a dramatic reduction from the standard 35 days through growth factor-based protocols^26^. Characterizations of TFiMGLs indicated that they possessed transcriptomic and functional resemblance to primary human microglia. We also explored the possibility of using single-cell atlas for guiding cell fate engineering by building a single-cell reference and mapping our pooled screen scRNA-seq data to it. We observed an increased percentage of microglia mapping cells from the second iteration when compared to the first. The high expression of key TFs like CEBPA and SPI1 in the microglia containing cluster also confirmed their ability to drive microglia differentiation from iPSCs. By doing genome wide regression analysis between TF barcode counts and gene expression levels, we revealed TF-gene regulatory relationships present in these pooled screens.

During this study, we noted several technological challenges that could be addressed in future studies. Most current TF-based differentiation protocols rely on a one-time induction of TF expression, while lacking the capability for sequential induction. Despite this, current strategies have successfully generated certain cell types, including multiple types of neurons^75^, endothelial cells^2^, and the induction of iPSC itself^76^. However, during development *in vivo* coordinated gene programs are sequentially activated, as observed in time-resolved transcriptomic analysis of developing tissues^77^. This feature could be re-created by identifying orthogonal induction system with a comparable strength to the doxycycline-inducible system or developing tunable gene circuits. With these tools, it will be possible to test whether sequential TF expression can lead to improved differentiation accuracy. In addition to temporal control of TF expression, the ability to regulate expression levels of individual TFs could also lead to improvements in differentiation. There were two manifestations of this pattern in the current study. In the first case, although CEBPA and FLI1 expressed by themselves led to cell death, the presence in the SFC cassette enabled cell survival and differentiation. The reduced expression levels of CEBPA and FLI1 likely and potential interactions with other TFs could also explain why we were able to observe an extensive GRN for CEBPA from the first pooled screen, which would not be possible due to toxicity in CEBPA-expressing cells. The second case can be observed in the effect that different sequential arrangements of TFs in the polycistronic cassettes led to different levels of downstream microglial protein expression. The effect of TF stoichiometry on differentiation efficiency has also been observed for cardiac myocyte programming^78^. While the positional effects in a polycistronic cassette offer one way to explore the stoichiometry space, development of new titratable promoters allowing turning of individual genes could be an important tool for cell fate engineering.

We demonstrated the potential for using primary single-cell atlas to guide iPSC differentiation efforts. We note that these approaches will improve with better quality atlases. A cell atlas with wide representation of human tissues, deep sequencing coverage, and reliable cellular-level annotation is ideal for guiding cell fate engineering. A number of methods for scRNA-seq datasets integration have been developed^79^, with the goal of enabling comparison between batches of data. However, because construction of a comprehensive cell atlas would need to integrate data from tens to hundreds of separately acquired datasets, the accuracy and computational efficiency of current integration strategies require improvement. We also note, that current reference mapping methods were designed to project new data from the same tissue to old datasets, or projecting data from the same tissue but acquired by different modalities. As a result, these methods are not optimized for iPSC-derived cells with incomplete conversions, leading to partially resemble multiple primary cells types. A reference mapping method that provides probability-based quantitative measurement and rejection options is needed to address this issue. Furthermore, we believe experimental strategies might be devised to improve mapping of differentiated iPSCs to single cell reference data sets. For example, spiking in a standard set of differentiated cell types could be used as landmarks to validate single cell profile mapping.

As demonstrated in this study, we used microglia as a model target to develop iterative screening methods for identifying TFs that drive iPSC differentiation towards a specific cell type. We found that the combination of SPI1, CEBPA, FLI1, MEF2C, CEBPB, and IRF8 could produce microglia-like cells from iPSCs in as quickly as four days. We built computationally a comprehensive single-cell reference atlas and used it to validate the results from our iterative screening. We also used a stepwise regression model to discover TF-gene regulatory relationship from the scRNA-seq data. We believe the TFiMGLs can be used as a model for human microglia and facilitate basic and translational research that would benefit from a rapid turnaround time. We also envisage that the methods of iterative pooled TF screen, single-cell reference atlas mapping, and TF-gene regulation analysis will find their utility in other cell type targets for iPSC differentiation.

## Acknowledgments

The authors would like to acknowledge Gabriel Filsinger, Yu Wang, Cory Smith, Dima Ter-Ovanesyan, Michael Chou for helpful discussions. The authors would also like to thank the Biopolymers Facility at HMS, Research Computing at HMS, Harvard Chan Bioinformatics Core, and HMS Immunology Flow Cytometry Core Facility for technical assistance. This work has been supported by the National Human Genome Research Institute of the National Institutes of Health [RM1 HG008525 to G.M.C.] and the Lipper Foundation. Some schematics in this paper were created using BioRender.com.

## Author contributions

G.M.C. and S.L. conceived the project. S.L., with guidance from A.H.M.N. and P.K., performed early exploratory experiments. S.L., G.M.C., L.L., F.Z. designed overall experimental and analytical strategies. S.L. performed experiments with help from B.S., Y.C., E.A., M.G-C., C-T.W., J.H., Y.T., and G.C. L.L. helped with single-cell TF barcode quantification and regression analysis. F.Z. and S.R. helped with cell atlas integration and mapping. J.A., J.T., E.L. provided significant discussion and input over project design. S.L. wrote the manuscript with help from L.L., and with input and feedback from all authors.

## Competing financial interests

G.M.C, P.K., and A.H.M.N. are co-founders of and have equity in GC Therapeutics, Inc. Full disclosure for GMC is available at arep.med.harvard.edu/gmc/tech.html.

## Code availability

Code used in this study for bulk RNA-seq, scRNA-seq, reference mapping, and regression analysis will be made available upon publication.

## Data availability

Bulk RNA-seq data, scRNA-seq data, and the single-cell reference atlas will be made available upon publication.

## Online Methods

### Barcoded TF expression vector construction

All TFs used in this study were obtained from the TFome collection^2^ in pDONR format. For expression in hiPSCs, a PiggyBac integrating Dox-inducible vector pBAN^2^ was used. To create barcoded pBAN expression vector (pBAN-BC), the original pBAN was digested with AgeI and KpnI, followed by ligation of a gBlock (IDT DNA) containing the same excised piece with an additional 20-bp random barcode. After bacteria transformation, individual colonies were expanded and extracted for plasmid DNA. Gateway cloning was used to transfer each TF from pDONR to pBAN-BC vector. Barcode sequence for each TF was confirmed by Sanger sequencing.

### Cell culture

hiPSCs were culture in mTeSR Plus media (Stemcell Technologies, 100-0276) on multi-wells plates coated with Matrigel (Corning, 354277) or Cultrex (Bio-Techne Corporation, 3434-005-02). For passaging, cells were dissociated with TrypLE Express (Life Technologies, 12604013) and seeded into fresh plate and media containing 10 μM Y-27632 ROCK inhibitor (Millipore, 688001) for 24 hours. Daily media change was performed until cells were ready for another passaging or downstream experiments.

### Nucleofection, TF integration and differentiation

TF (pBAN-TF-BC) and Super PiggyBac (SPB) Transposase (System Biosciences, PB210PA-1) expression vectors were mixed at a mass ration of 4:1 and transfected into hiPSCs using P3 Primary Cell 4D-Nucleofector X Kit L (Lonza, V4XP-3024) on a 4D-Nucleofector X Unit (Lonza, AAF-1002X) following manufacturer’s instructions. For the two pooled TF screenings, 600,000 cells were transfected with 5 μg of DNA and seeded into one well of a 6-well plate. For individual TF combinations, 120,000 cells were transfected with 2.5 μg of DNA and seeded into one well of a 12-well plate. Program CB150 was used for the nucleofections. 48 hours after nucleofection, 1 μg/mL of puromycin (Gibco, A1113803) or 50 μg/mL of zeocin (Gibco, R25001) was added to the culture for the selection of TF-integrated cells. Cells were passaged again when reaching 80% confluency. For induction of TF expression, cells were seeded into mTeSR Plus media containing 0.5 μg/mL doxycycline (Sigma-Aldrich, D3072) and 10 μM Y-27632 ROCK inhibitor and were changed into media only containing doxycycline after 24 hours.

### Flow cytometry and sorting

For cytometry analysis, cells were dissociated with TrypLE Express for 5 minutes at 37 degree, diluted with twice the volume of Cell Staining Buffer (Biolegend, 420201) and centrifuged at 200 g for 3 minutes to remove the digesting enzyme. Cells were then incubated with 25 μg/mL of Human Fc Block (BD Biosciences, 564219) diluted in Cell Staining Buffer for 15 minutes on ice, followed immediately by staining with fluorescently conjugated antibodies or isotype controls at proper dilution for 30 minutes on ice. Antibodies were diluted in Cell Staining Buffer and Human Fc Block was not removed from the mixture. After antibody staining, cells were washed twice with Cell Staining Buffer before being put through 35 μm nylon mesh into a 5 mL round bottom polystyrene tube (Falcon, 352235). Flow cytometry data was acquired on a BD LSRFortessa Cell Analyzer. For cell sorting, the staining protocol was the same except for that Cell Staining Buffer was replaced with mTeSR Plus media in order to maintain the best viability of cells. Cell sorting was performed on a BD FACSAria Cell Sorter. Flow cytometry antibodies used in this study were: FITC-TRA-1-60 (BD Biosciences, 560380), BV421-CX3CR1 (Biolegend, 341620), PE-P2RY12 (Biolegend, 392104), APC-CD11b (Biolegend, 101212). Isotype controls used were: BV421-Rat IgG2b (Biolegend, 400640), PE-Mouse IgG2a (Biolegend, 400214), APC-Rat IgG2b (Biolegend, 400612).

### scRNA-seq library preparation

scRNA-seq experiments were performed using 10x Genomics Chromium Single Cell 3’ Reagent Kits v3 or v3.1 following the manufacturer’s instruction. 5000 single cells were calculated as targeted input for each sample. For the first iteration, 10% of stem cells were spiked in as undifferentiated control. For the second iteration, 5% stem cells and 5% MG3.1-SFC were spiked in as undifferentiated and initial differentiation control. The only modification made to the protocol was at the Sample Index PCR step, where 5 μL of the PCR mix was taken out and mixed with 0.5 μL 1000X SYBR Gold (Invitrogen, S11494) for a qPCR reaction. The optimal amplification cycle was determined as the cycle just before half maximum of the total signal. Final libraries were sequenced on NextSeq 500 or NovaSeq with a goal of at least 30,000 reads per cell.

### TF barcode amplicon library preparation

Because after the cDNA amplification step in the 10x scRNA-seq protocol the amplicons contained cell barcodes, UMIs, and TF barcodes, these cDNAs could be used as the template for further amplification of TF-cell barcodes. Two sequential PCR reactions were performed, each was accompanied by a SYBR Gold spike-in qPCR to determine the optimal cycle number as described in “scRNA-seq library preparation”. For PCR1, NGS10x-F-i7-BC-PCR1F and i5000 were used as primers. A 50 μL PCR1 reaction contains 25 μL Q5 Hot Start High-Fidelity 2X Master Mix (New England Biolabs, M0494L), 5 μL amplified cDNA, 2.5 μL of both primers at 10 μM stock concentration, and 15 μL nuclease-free water. PCR1 program was initial denaturation, 98 degrees, 30 seconds; 11-13 cycles (qPCR determined) of 98 degrees, 10 seconds, 67 degrees, 30 seconds, 72 degrees, 30 seconds; final extension, 72 degrees, 2 minutes. PCR1 reaction was purified with 1.2X SPRIselect beads (Beckman Coulter, B23318) following standard protocol. The sample was eluted in 20 μL water. For PCR2, i7000, P5, and P7 were used as primers. A 50 μL PCR2 reaction contains 25 μL Q5 Hot Start High-Fidelity 2X Master Mix, 10 μL PCR1 product, 2.5 μL of all three primers at 10 μM stock concentration, and 7.5 μL nuclease-free water. PCR2 program was initial denaturation, 98 degrees, 30 seconds; 4-5 cycles (qPCR determined) of 98 degrees, 10 seconds, 67 degrees, 30 seconds, 72 degrees, 30 seconds; final extension, 72 degrees, 2 minutes. PCR2 product was purified the same as PCR1. Final libraries were submitted for MiSeq v3 with paired-end reads of 80 cycles from either direction.

Primer sequences:

NGS10x-F-i7-BC-PCR1F: GGAGTTCAGACGTGTGCTCTTCCGATCTCTTTTCCAAGCACCTGCTACATAG

i5000:

AATGATACGGCGACCACCGAGATCTACACaactcgctACACTCTTTCCCTACACGACGCTCTTC CGATCT (lower case region represents a sample-specific barcode)

i7000:

CAAGCAGAAGACGGCATACGAGATtcgccttaGTGACTGGAGTTCAGACGTGTGCTCTTCCGA TCT (lower case region represents a sample-specific barcode)

P5: AATGATACGGCGACCACCGA

P7: CAAGCAGAAGACGGCATACGA

### Analysis of scRNA-seq and TF barcode-seq data

For scRNA-seq, raw FASTQ files were aligned to GRCh38 and quantified using Cell Ranger. Detailed information about cell number, read depth and gene detected is visualized in **Supplementary Figure S2** and **S6**. Seurat was used to performed cell filtering, data normalization and clustering. The generated Seurat object also contained the single-cell raw expression matrix for all genes. For TF barcode-seq, in the paired-end MiSeq data, one of the read pair contains the 20 bp TF barcode while the other one contains the 16 bp cell barcode and the 12 bp UMI. By matching the names of the reads within the pair, three sequences were compiled into one table with three columns: TF-BC, cell-BC, UMI. To remove duplicated reads from the same molecule, duplicated rows that has the same value I for all three columns were removed. Then the table was counted and reshaped into a frequency table where the row names represent cell and column names represent TF. This table contains the raw counts of each TF barcode in all single cells. Because the TF barcodes were amplified from the cDNA during library preparation, we normalized the TF barcode count with the number of total RNA UMIs detected in each cell, reasoning that cells with more total UMIs were likely to have more reads for TF barcode. The raw gene expression matrix and normalized TF count matrix were used to identify which TF barcodes were likely to induce microglial gene expression. Specifically, the expression of microglial genes was binarized, with any cell had a non-zero expression being 1. Then between the two groups of cells 0 or 1 microglial gene expression, a Wilcoxon rank sum test was performed for all barcoded TFs to determine which TF(s) had a higher * expression in cells expressing microglial genes. The TFs were ranked by -log_10_(p-value).

### Bulk RNA-seq library preparation

Cultured cells were dissolved directly with TRIzol (Thermo Fisher Scientific, 15596018) for total RNA purification with Direct-zol RNA MiniPrep Kit (Zymo Research, R2050). RNA concentration was quantified with Qubit RNA HS Assay Kit (Thermo Fisher Scientific, Q32852). RNA integrity was confirmed by presence of 18S and 28S bands on a 2% E-Gel EX Agarose Gel (Thermo Fisher Scientific, G402002). Between 100 ng to 1000 ng total RNA was used as input for mRNA enrichment using NEBNext Poly(A) mRNA Magnetic Isolation Module (New England Biolabs, E7490), followed by library construction with NEBNext Ultra II Directional RNA Library Prep Kit (New England Biolabs, E7760S) following the manufacturer’s instructions. Biopolymers Facility at Harvard Medical School performed library QC and sequencing.

### Analysis of bulk RNA-seq data

For both in-house generated sample and datasets downloaded from GEO, raw FASTQ files were aligned to GRCh38 and quantified using STAR 2.5.2b. Regularized-logarithm (rlog) transformation was applied to the raw counts before visualization using PCA. For analysis where data from multiple sources were involved, ComBat-seq was used for batch correction before PCA. Differential gene expression analysis was conducted with DESeq2^47^. Pathway enrichment and GSEA analysis were performed with clusterProfiler^80^.

### Immunofluorescence (IF)

IF experiments were performed in μ-Plate 96 Well Black plate (ibidi, 89626). After media removal, cells were fixed with 4% paraformaldehyde (Electron Microscopy Sciences, 15710) in 1x phosphate buffered saline (PBS) (Thermo Fisher Scientific, 10010072) for 15 minutes at room temperature (RT). Cells were rinsed three times with PBS before proceeding to permeabilization or blocking. For staining of Oct-3/4 and PU.1, cells were permeabilized, while not for cell surface proteins’ staining. Permeabilization was conducted with 0.25% Triton-X-100 (Thermo Fisher Scientific, 85111) in 1x PBS for 15 minutes at RT followed by three rinses with PBS. Cells were then blocked with 1% bovine serum albumin (BSA) in PBS for one hour at RT. For primary and secondary antibody staining, antibodies were diluted in PBS with 1% BSA and incubated with cells for one hour at RT. Three 5-minute washes with PBS were used to remove excessive antibodies after staining. Cells were directly imaged in plate on a Nikon Ti2 Eclipse inverted microscope with a Plan Apo Lambda DM 60× (1.4 NA, Ph3) oil objective and an Andor Zyla sCMOS camera. Images were acquired by NIS-Element AR software. All antibodies were used at 1:200 dilution. Primary IF antibodies used in this study were: Oct-3/4 (Santa Cruz Biotechnology, sc-5279), PU.1 (Thermo Fisher Scientific, PA5-17505), CD11b (BioLegend, 101202), P2RY12 (Thermo Fisher Scientific, 702516), CX3CR1 (Abcam, ab8021).

### Phagocytosis assay

Differentiated cells were incubated with 20 μg/mL of pHrodo Green S. aureus BioParticles (Thermo Fisher Scientific, P35382) for 0-4 hours in mTeSR Plus media in the presence of 100 μg/ml Penicillin-Streptomycin (Corning, 30-002-CI). After removal of excessive particles with PBS washes, cells were harvested for antibody (CX3CR1, P2RY12, CD11b) staining and flow cytometry analysis as described in previous section. Remaining stained cells after flow cytometry was transferred into μ-Plate 96 Well Black plate for fluorescence microscopy to confirm the intracellular localization of the particles. This step needs to be conducted swiftly after flow cytometry in order to avoid changing of cellular morphology due to cell death.

### Calcium imaging

Calcium imaging experiment was conducted in standard 12-well cell culture plates. Differentiated cells were incubated with 1 μg/mL Fluo-4 AM calcium indicator (Thermo Fisher Scientific, F23917) in 1 mL of mTeSR Plus media for 30 minutes in a cell culture incubator. Excessive dye was washed away with two 1 mL media washes. After adding 1 mL of fresh mTeSR Plus, the cells were put on stage in a microscope inside the incubator. Images acquisition started without stimulation for 90 seconds to determine baseline signal. One image was acquired every three seconds, the fastest possible on the instrument. After 90 seconds 1 mL of media containing 150 μM ADP was added to the cells while imaging was continuing. The total length of imaging was 10 minutes. Fluorescent signal was quantified and plotted in MATLAB.

### Amyloid fibrillation

Aβ fibrillation experiments were performed using SensoLyte Thioflavin T β-Amyloid (1-42) Aggregation Kit (AnaSpec, AS-72214) according to manufacturer’s instruction. The reaction was set up in μ-Plate 96 Well Black plate. Data was acquired on a plate reader with excitation/emission = 440 nm/484 nm at 37 degree once every 5 minutes for 3 hours.

### Preparation of datasets for building single-cell reference atlas

Files containing raw counts of 10x Genomics Chromium scRNA-seq data for different human tissues were download from PanglaoDB (https://panglaodb.se/index.html). Human primary brain single-cell data from gestational weeks 6-22 were downloaded from Organoid Report Card (https://cells.ucsc.edu/?ds=organoidreportcard). Human endometrium single-cell data were download from GEO GSE111976. All sample went through manual cell filtering using Seurat with different filters on number of gene, UMI, and percentage of mitochondria genes (**Supplementary Table S4**). Tissue annotation was compiled through manual curation of each study by checking what tissue/cell types were used (**Supplementary Table S3**). All raw counts were merged into one sparse matrix, which was then used as input for data integration.

### Single-cell atlas integration and mapping

Data integration and projection using Harmony and Symphony was carried out following instructions from the authors on GitHub (https://github.com/immunogenomics/harmony; https://github.com/immunogenomics/symphony). For Harmony integration, RunHarmony function was used. Parameters different from default settings were epsilon.cluster=-Inf, epsilon.harmony=-Inf. Batch correction were performed based on tissue types labeld in Figure 5a. For reference mapping with Symphony, buildReferenceFromSeurat function was used to create the reference object, and mapQuery function was used to map iPSC-derived cells to the reference atlas. Due to the size of the data, these steps were performed on the O2 cluster of Harvard Medical School with at least 180 Gb memory and 8 cores. Most R objects along the pipeline could be saved as standard R files for repeated analysis, except for the UMAP model file, which required saving and loading through the “uwot” package^81^. Code used for integration and projection, together with key reference and annotations files that could be of use for future explorations are shared along with this manuscript.

### TF-gene stepwise regression model construction

The stepwise regression (or stepwise selection) is a regression model that iteratively adds and removes predictors in the predictive model to find the subset of variables in the data set resulting in the best performance, and consequently lowering the perdition error in the model. During the process, the value of the statistical test is used to screen the variables. If the value is less than or equal to 0.05, then the variable enters the regression model, and the selected variable is the independent variable of the regression model. For the construction of the model:

Step 1: Establish *P* regression models between the independent variables *X*_1_, *X*_2_,…,*X_p_* (*number = P*) and the dependent variable *Y* respectively,

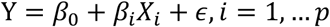 Calculate the statistical value of the F-test with the regression coefficient 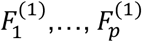, and take the maximum value 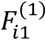,

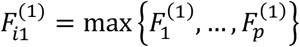 For a given significance level *α*, the threshold value is *F*^1^. If 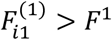, then *X*_*i*1_ will be included in the regression model and recorded as the set of selected variable indicators as *I*_1_.
Step 2: Establish a binary regression model of the dependent variable *Y* and the independent variable subset {*X*_*i*1_,…,*X*_1_}, {*X*_*i*_,…, *X*_*i*1–1_}, {*X*_*i*_,…, *X*_*i*1+1_}, calculate the statistical value of the F-test with the regression coefficient 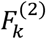 and take the maximum value 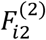,

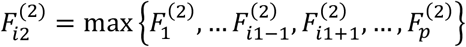 For a given significance level *α*, record the corresponding critical value as *F*^(2)^. If 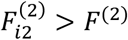, then the variable is introduced into the regression model. Otherwise, the variable introduction process is terminated.
Step 3: Repeat Step 2 with the subset of variables {*X*_*i*1_, *X*_*i*2_, *X_k_*}. This step is repeated by selecting an independent variable that is not introduced into the regression model until the test does not introduce any variables.

### TF-gene network visualization

Both p-values and coefficients in the regression analysis work together to represent relationships in the model about the significant factors. The coefficients describe the mathematical relationship between each independent variable and the dependent variable. The p-values for the coefficients indicate whether these relationships are statistically significant. We selected the 250 TF-gene combinations from the first pooled screen and 422 in the second with the criteria Abs(coefficient) > 0.1 & -log_10_(p-value) > 20, then visualized them in GEPHI. Nodes are the genes and TFs, edges are the regression coefficients (an unstandardized effect size because they indicate the strength of the relationship between variables), and colors are based on the modularity from the network module algorithm^82^.

## References

1. Regev, A., Teichmann, S. A., Lander, E. S., Amit, I. & Benoist, C. Science forum: the human cell atlas. elife (2017).

2. Ng, A. H. M. et al. A comprehensive library of human transcription factors for cell fate engineering. Nat. Biotechnol. 39, 510–519 (2021).

3. Ginhoux, F. et al. Fate mapping analysis reveals that adult microglia derive from primitive macrophages. Science 330, 841–845 (2010).

4. Hoeffel, G. et al. C-Myb(+) erythro-myeloid progenitor-derived fetal monocytes give rise to adult tissue-resident macrophages. Immunity 42, 665–678 (2015).

5. Crotti, A. & Ransohoff, R. M. Microglial Physiology and Pathophysiology: Insights from Genome-wide Transcriptional Profiling. Immunity 44, 505–515 (2016).

6. Nayak, D., Roth, T. L. & McGavern, D. B. Microglia development and function. Annu. Rev. Immunol. 32, 367–402 (2014).

7. Nimmerjahn, A., Kirchhoff, F. & Helmchen, F. Resting microglial cells are highly dynamic surveillants of brain parenchyma in vivo. Science 308, 1314–1318 (2005).

8. Matcovitch, O. Microglia development follows a stepwise program to regulate brain homeostasis. Natan

9. Salter, M. W. & Beggs, S. Sublime microglia: expanding roles for the guardians of the CNS. Cell 158, 15–24 (2014).

10. Colonna, M. & Butovsky, O. Microglia function in the central nervous system during health and neurodegeneration. Annu. Rev. Immunol. 35, 441–468 (2017).

11. Salter, M. W. & Stevens, B. Microglia emerge as central players in brain disease. Nat. Med. 23, 1018–1027 (2017).

12. Mathys, H. et al. Single-cell transcriptomic analysis of Alzheimer’s disease. Nature 570, 332–337 (2019).

13. Keren-Shaul, H. et al. A Unique Microglia Type Associated with Restricting Development of Alzheimer’s Disease. Cell 169, 1276–1290.e17 (2017).

14. Yeh, F. L., Hansen, D. V. & Sheng, M. TREM2, microglia, and neurodegenerative diseases. Trends Mol. Med. 23, 512–533 (2017).

15. Ulland, T. K. et al. TREM2 maintains microglial metabolic fitness in alzheimer’s disease. Cell 170, 649–663.e13 (2017).

16. Dello Russo, C. et al. The human microglial HMC3 cell line: where do we stand? A systematic literature review. J. Neuroinflammation 15, 259 (2018).

17. Timmerman, R., Burm, S. M. & Bajramovic, J. J. An Overview of in vitro Methods to Study Microglia. Front. Cell Neurosci. 12, 242 (2018).

18. Smith, A. M. & Dragunow, M. The human side of microglia. Trends Neurosci. 37, 125–135 (2014).

19. Watkins, L. R. & Hutchinson, M. R. A concern on comparing’apples’ and’oranges’ when differences between microglia used in human and rodent studies go far, far beyond simply species …. Trends in neurosciences (2014).

20. Muffat, J. et al. Efficient derivation of microglia-like cells from human pluripotent stem cells. Nat. Med. 22, 1358–1367 (2016).

21. Abud, E. M. et al. iPSC-Derived Human Microglia-like Cells to Study Neurological Diseases. Neuron 94, 278–293.e9 (2017).

22. Pandya, H. et al. Differentiation of human and murine induced pluripotent stem cells to microglia-like cells. Nat. Neurosci. 20, 753–759 (2017).

23. Haenseler, W. et al. A Highly Efficient Human Pluripotent Stem Cell Microglia Model Displays a Neuronal-Co-culture-Specific Expression Profile and Inflammatory Response. Stem Cell Rep. 8, 1727–1742 (2017).

24. Douvaras, P. et al. Directed differentiation of human pluripotent stem cells to microglia. Stem Cell Rep. 8, 1516–1524 (2017).

25. Takata, K. et al. Induced-Pluripotent-Stem-Cell-Derived Primitive Macrophages Provide a Platform for Modeling Tissue-Resident Macrophage Differentiation and Function. Immunity 47, 183–198.e6 (2017).

26. McQuade, A. et al. Development and validation of a simplified method to generate human microglia from pluripotent stem cells. Mol. Neurodegener. 13, 67 (2018).

27. Speicher, A. M., Wiendl, H., Meuth, S. G. & Pawlowski, M. Generating microglia from human pluripotent stem cells: novel in vitro models for the study of neurodegeneration. Mol. Neurodegener. 14, 46 (2019).

28. Xu, R. et al. Human iPSC-derived mature microglia retain their identity and functionally integrate in the chimeric mouse brain. Nat. Commun. 11, 1577 (2020).

29. Vierbuchen, T. et al. Direct conversion of fibroblasts to functional neurons by defined factors. Nature 463, 1035–1041 (2010).

30. Tsunemoto, R. et al. Diverse reprogramming codes for neuronal identity. Nature 557, 375–380 (2018).

31. Busskamp, V. et al. Rapid neurogenesis through transcriptional activation in human stem cells. Mol. Syst. Biol. 10, 760 (2014).

32. Kierdorf, K. et al. Microglia emerge from erythromyeloid precursors via Pu.1- and Irf8-dependent pathways. Nat. Neurosci. 16, 273–280 (2013).

33. Smith, A. M. et al. The transcription factor PU.1 is critical for viability and function of human brain microglia. Glia 61, 929–942 (2013).

34. Gosselin, D. et al. An environment-dependent transcriptional network specifies human microglia identity. Science 356, (2017).

35. Galatro, T. F. et al. Transcriptomic analysis of purified human cortical microglia reveals age-associated changes. Nat. Neurosci. 20, 1162–1171 (2017).

36. Zhong, S. et al. A single-cell RNA-seq survey of the developmental landscape of the human prefrontal cortex. Nature 555, 524–528 (2018).

37. Olah, M. et al. A transcriptomic atlas of aged human microglia. Nat. Commun. 9, 539 (2018).

38. Butovsky, O. et al. Identification of a unique TGF-β-dependent molecular and functional signature in microglia. Nat. Neurosci. 17, 131–143 (2014).

39. Wehrspaun, C. C., Haerty, W. & Ponting, C. P. Microglia recapitulate a hematopoietic master regulator network in the aging human frontal cortex. Neurobiol. Aging 36, 2443.e9–2443.e20 (2015).

40. Avellino, R. & Delwel, R. Expression and regulation of C/EBPα in normal myelopoiesis and in malignant transformation. Blood 129, 2083–2091 (2017).

41. Lichtinger, M. et al. RUNX1 reshapes the epigenetic landscape at the onset of haematopoiesis. EMBO J. 31, 4318–4333 (2012).

42. Starck, J. et al. Spi-1/PU.1 is a positive regulator of the Fli-1 gene involved in inhibition of erythroid differentiation in friend erythroleukemic cell lines. Mol. Cell. Biol. 19, 121–135 (1999).

43. Hoeffel, G. & Ginhoux, F. Ontogeny of Tissue-Resident Macrophages. Front. Immunol. 6, 486 (2015).

44. Bazan, J. F. et al. A new class of membrane-bound chemokine with a CX3C motif. Nature 385, 640–644 (1997).

45. Hughes, P. M., Botham, M. S., Frentzel, S., Mir, A. & Perry, V. H. Expression of fractalkine (CX3CL1) and its receptor, CX3CR1, during acute and chronic inflammation in the rodent CNS. Glia 37, 314–327 (2002).

46. Liu, Z. et al. Systematic comparison of 2A peptides for cloning multi-genes in a polycistronic vector. Sci. Rep. 7, 2193 (2017).

47. Love, M. I., Huber, W. & Anders, S. Moderated estimation of fold change and dispersion for RNA-seq data with DESeq2. Genome Biol. 15, 550 (2014).

48. Liberzon, A. et al. The Molecular Signatures Database (MSigDB) hallmark gene set collection. Cell Syst. 1, 417–425 (2015).

49. Cahan, P. et al. CellNet: network biology applied to stem cell engineering. Cell 158, 903–915 (2014).

50. Kracht, L. et al. Human fetal microglia acquire homeostatic immune-sensing properties early in development. Science 369, 530–537 (2020).

51. Inoue, K. Purinergic systems in microglia. Cell Mol. Life Sci. 65, 3074–3080 (2008).

52. Di Virgilio, F., Ceruti, S., Bramanti, P. & Abbracchio, M. P. Purinergic signalling in inflammation of the central nervous system. Trends Neurosci. 32, 79–87 (2009).

53. Zhang, Y., Parmigiani, G. & Johnson, W. E. ComBat-seq: batch effect adjustment for RNA-seq count data. NAR Genom. Bioinform. 2, lqaa078 (2020).

54. Subramanian, A. et al. Gene set enrichment analysis: a knowledge-based approach for interpreting genome-wide expression profiles. Proc. Natl. Acad. Sci. USA 102, 15545–15550 (2005).

55. Ivashkiv, L. B. IFNγ: signalling, epigenetics and roles in immunity, metabolism, disease and cancer immunotherapy. Nat. Rev. Immunol. 18, 545–558 (2018).

56. Zhong, L. et al. Amyloid-beta modulates microglial responses by binding to the triggering receptor expressed on myeloid cells 2 (TREM2). Mol. Neurodegener. 13, 15 (2018).

57. Zhao, W. et al. TDP-43 activates microglia through NF-κB and NLRP3 inflammasome. Exp. Neurol. 273, 24–35 (2015).

58. Zhang, F. et al. IFN-γ and TNF-α drive a CXCL10+ CCL2+ macrophage phenotype expanded in severe COVID-19 lungs and inflammatory diseases with tissue inflammation. Genome Med. 13, 64 (2021).

59. Franzén, O., Gan, L.-M. & Björkegren, J. L. M. PanglaoDB: a web server for exploration of mouse and human single-cell RNA sequencing data. Database (Oxford) 2019, (2019).

60. Bhaduri, A. et al. Cell stress in cortical organoids impairs molecular subtype specification. Nature 578, 142–148 (2020).

61. Wang, W. et al. Single-cell transcriptomic atlas of the human endometrium during the menstrual cycle. Nat. Med. 26, 1644–1653 (2020).

62. Korsunsky, I. et al. Fast, sensitive and accurate integration of single-cell data with Harmony. Nat. Methods 16, 1289–1296 (2019).

63. Kang, J. B. et al. Efficient and precise single-cell reference atlas mapping with Symphony. Nat. Commun. 12, 5890 (2021).

64. Van de Sande, B. et al. A scalable SCENIC workflow for single-cell gene regulatory network analysis. Nat. Protoc. 15, 2247–2276 (2020).

65. Hecker, M., Lambeck, S., Toepfer, S., van Someren, E. & Guthke, R. Gene regulatory network inference: data integration in dynamic models-a review. Biosystems 96, 86–103 (2009).

66. Matys, V. et al. TRANSFAC and its module TRANSCompel: transcriptional gene regulation in eukaryotes. Nucleic Acids Res. 34, D108–10 (2006).

67. Castro-Mondragon, J. A. et al. JASPAR 2022: the 9th release of the open-access database of transcription factor binding profiles. Nucleic Acids Res. 50, D165–D173 (2022).

68. Kanehisa, M., Furumichi, M., Sato, Y., Ishiguro-Watanabe, M. & Tanabe, M. KEGG: integrating viruses and cellular organisms. Nucleic Acids Res. 49, D545–D551 (2021).

69. Liu, Z.-P., Wu, C., Miao, H. & Wu, H. RegNetwork: an integrated database of transcriptional and post-transcriptional regulatory networks in human and mouse. Database (Oxford) 2015, (2015).

70. Oughtred, R. et al. The BioGRID database: A comprehensive biomedical resource of curated protein, genetic, and chemical interactions. Protein Sci. 30, 187–200 (2021).

71. Szklarczyk, D. et al. The STRING database in 2021: customizable protein-protein networks, and functional characterization of user-uploaded gene/measurement sets. Nucleic Acids Res. 49, D605–D612 (2021).

72. Maeda, T., Sugita, S., Kurimoto, Y. & Takahashi, M. Trends of Stem Cell Therapies in Age-Related Macular Degeneration. J Clin Med 10, (2021).

73. de Klerk, E. & Hebrok, M. Stem Cell-Based Clinical Trials for Diabetes Mellitus. Front. Endocrinol. (Lausanne) 12, 631463 (2021).

74. Shi, Y., Inoue, H., Wu, J. C. & Yamanaka, S. Induced pluripotent stem cell technology: a decade of progress. Nat. Rev. Drug Discov. 16, 115–130 (2017).

75. Flitsch, L. J., Laupman, K. E. & Brüstle, O. Transcription Factor-Based Fate Specification and Forward Programming for Neural Regeneration. Front. Cell Neurosci. 14, 121 (2020).

76. Takahashi, K. & Yamanaka, S. Induction of pluripotent stem cells from mouse embryonic and adult fibroblast cultures by defined factors. Cell 126, 663–676 (2006).

77. Haniffa, M. et al. A roadmap for the Human Developmental Cell Atlas. Nature 597, 196–205 (2021).

78. Wang, L. et al. Stoichiometry of Gata4, Mef2c, and Tbx5 influences the efficiency and quality of induced cardiac myocyte reprogramming. Circ. Res. 116, 237–244 (2015).

79. Tran, H. T. N. et al. A benchmark of batch-effect correction methods for single-cell RNA sequencing data. Genome Biol. 21, 12 (2020).

80. Wu, T. et al. clusterProfiler 4.0: A universal enrichment tool for interpreting omics data. Innovation (N Y) 2, 100141 (2021).

81. McInnes, L., Healy, J. & Melville, J. UMAP: Uniform Manifold Approximation and Projection for Dimension Reduction. arXiv (2018). doi:10.48550/arxiv.1802.03426

82. Blondel, V. D., Guillaume, J.-L., Lambiotte, R. & Lefebvre, E. Fast unfolding of communities in large networks. J. Stat. Mech. 2008, P10008 (2008).

