## Supplementary Figures for "Iterative transcription factor screening enables rapid generation of microglia-like cells from human iPSC"

**This supplementary file includes:**

Supplementary Figure S1-S17

**This supplementary file does not include:**

Supplementary Table S1-S6, which is included as a separate Excel file.

Supplementary Video S1-S3, which are included as separate MOV files.

CGCCTGGAGGAGGCCGTGTGGAGGCCCTACTACCCAGCTTTCTTGTACAAAGTGG
TTGCAGGAAAGCCAATCCCAAACCCACTCCTAGGTCTCGACAGTACAGCATAGAC
CGGTCCACCACCACCACCACCACTAAGGATCCGGGGTTGGGGTTGCGCCTTTTCC
AAGCACCTGCTACATAGCNNNNNNNNNNNNNNNNNNNNGGTACCCCTCGACTGTG
CCTTCTAGTTGCCAGCCATCTGTTGTTTGCCCCTCCCCCGTGCCT

End of TF DNA

attB2

V5 tag

Stop codon

6xHis tag

20 base barcodes

bGH poly(A) signal

Supplementary Figure S1. Sequence organization around 20 nt TF barcodes.

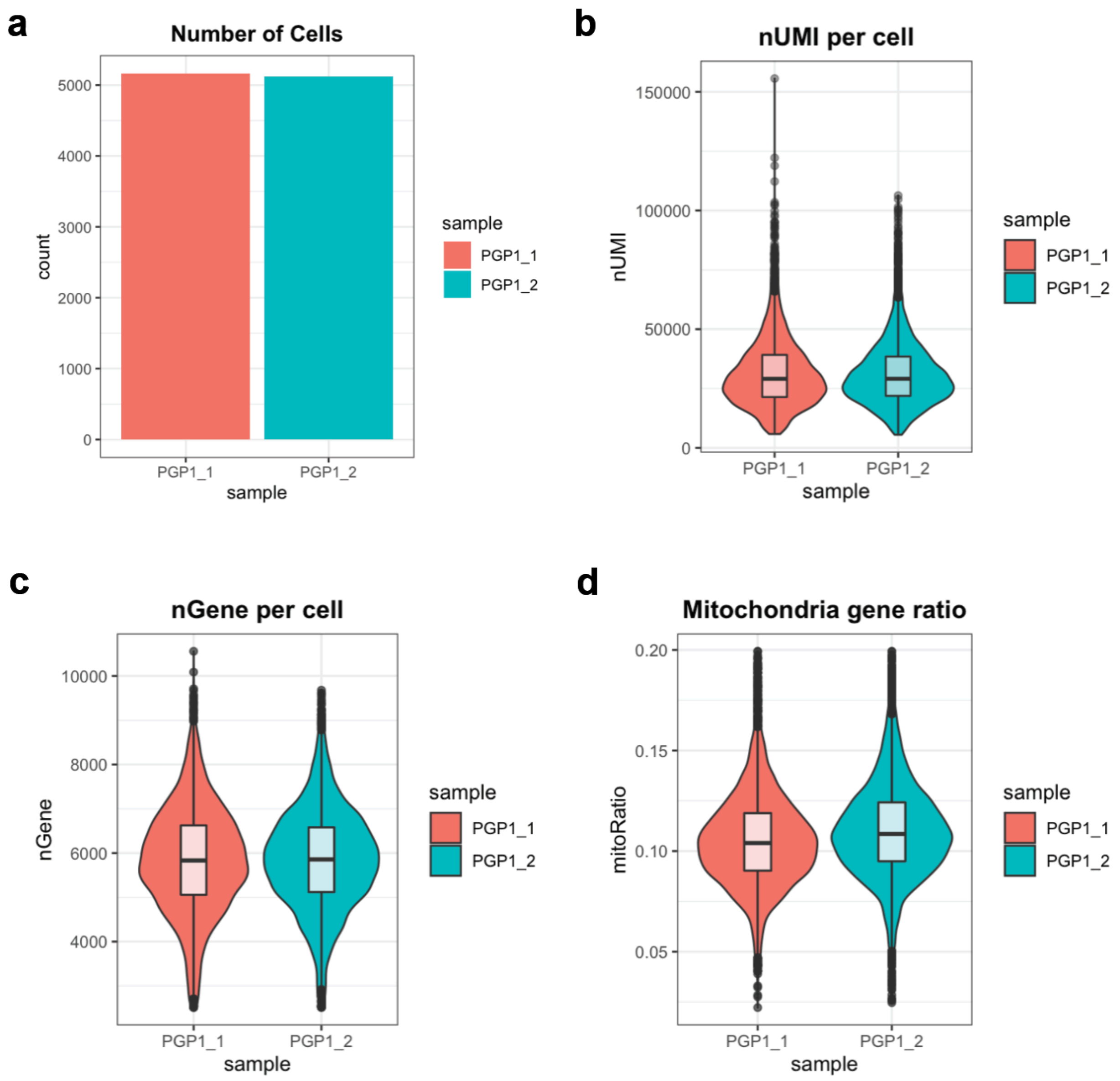


Supplementary Figure S2. Basic metric of scRNA-seq data from the first pooled screen of two samples (PGP1_1 and PGP1_2).

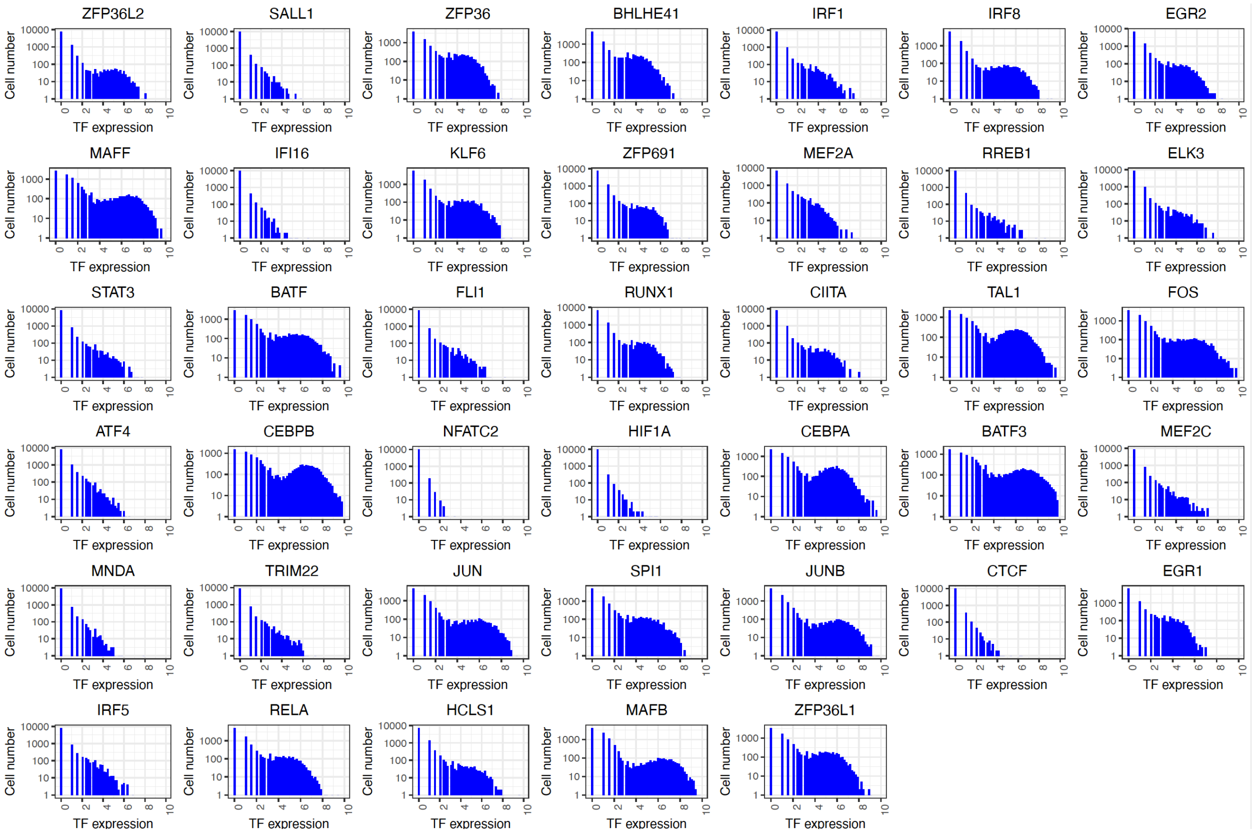


Supplementary Figure S3. Expression of the 40 TF barcodes in all single cells from the first pooled screen.

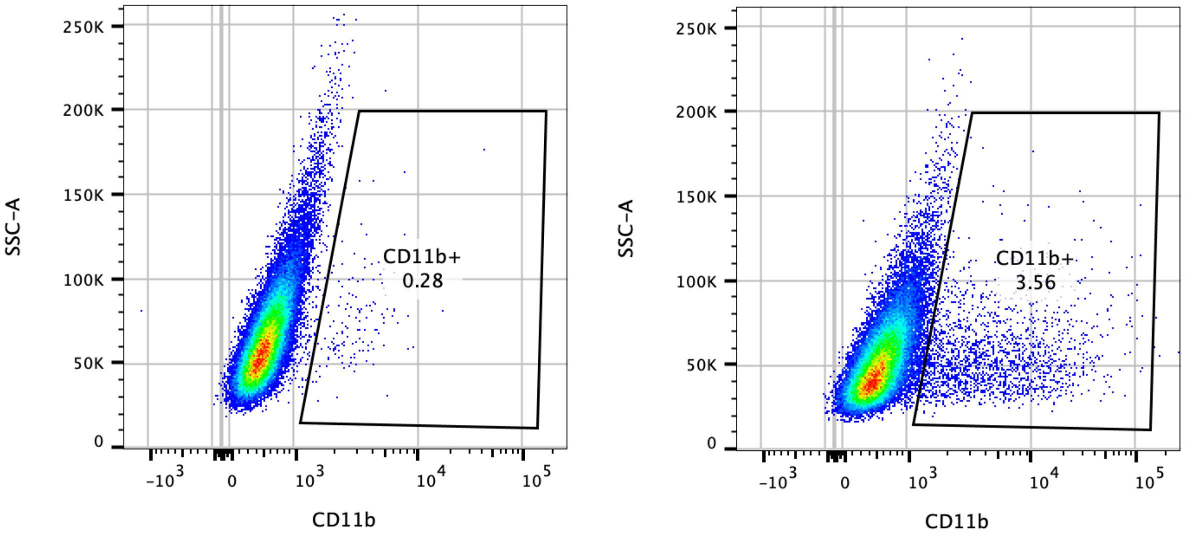


Supplementary Figure S4. Induction of CD11b expression by SPI1 along in hiPSCs.

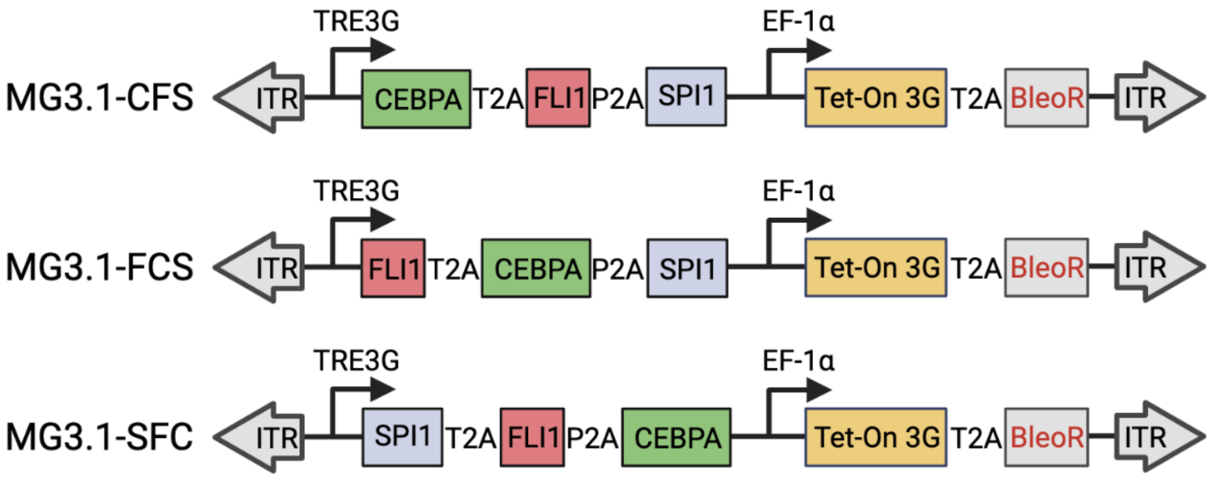


Supplementary Figure S5. Organization and naming of polycistronic cassettes for expression of CEBPA, FLI1 and SPI1.

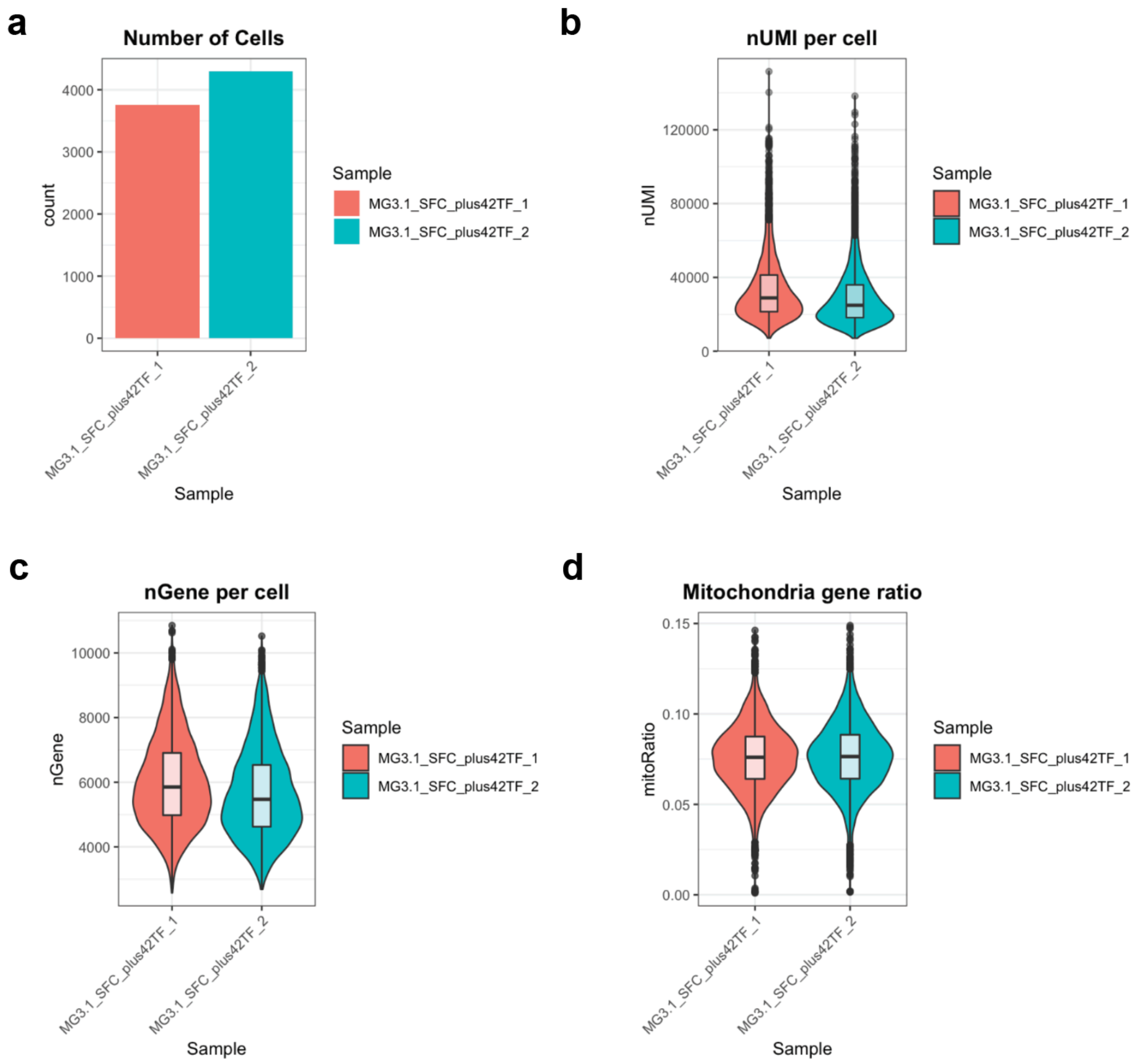


Supplementary Figure S6. Basic metric of scRNA-seq data from the second pooled screen of two samples (MG3.1_SFC_plus42TF_1 and (MG3.1_SFC_plus42TF_2).

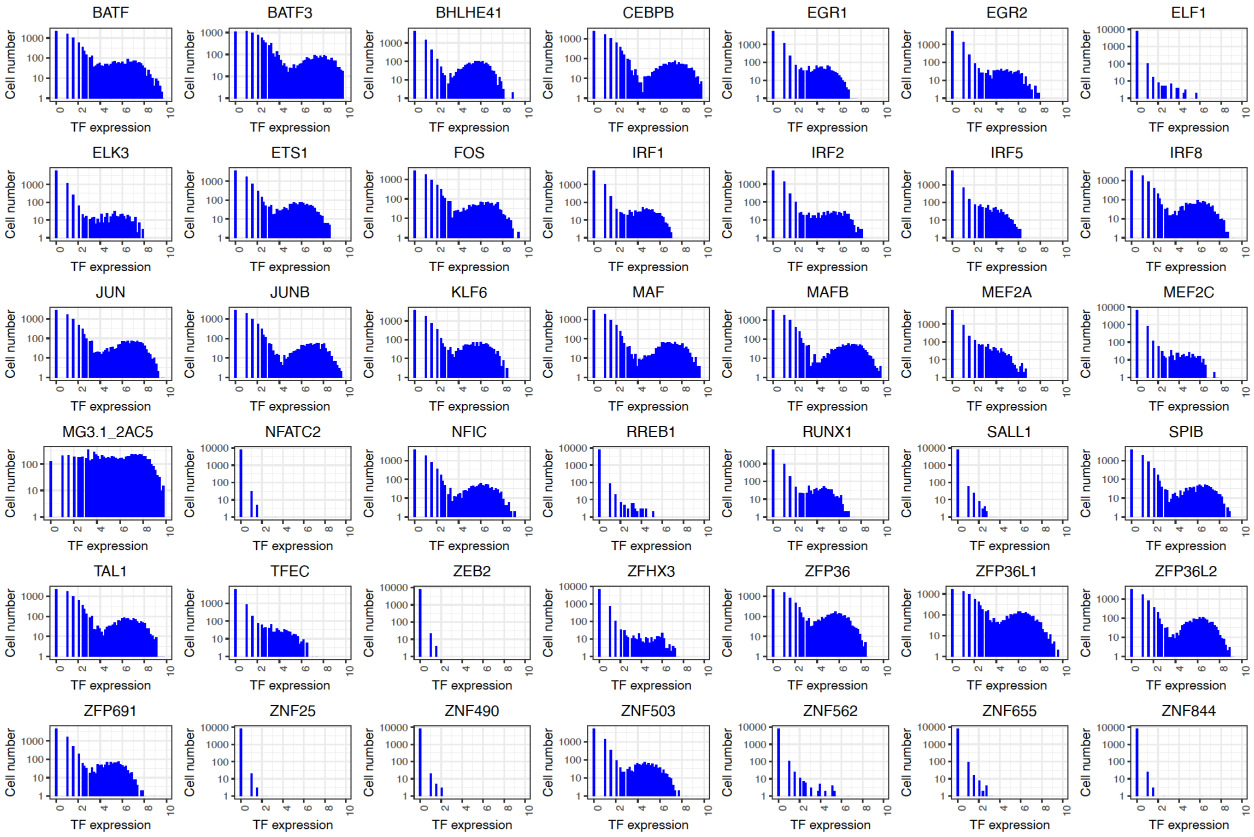


Supplementary Figure S7. Expression of the 42 TF barcodes in all single cells from the second pooled screen.

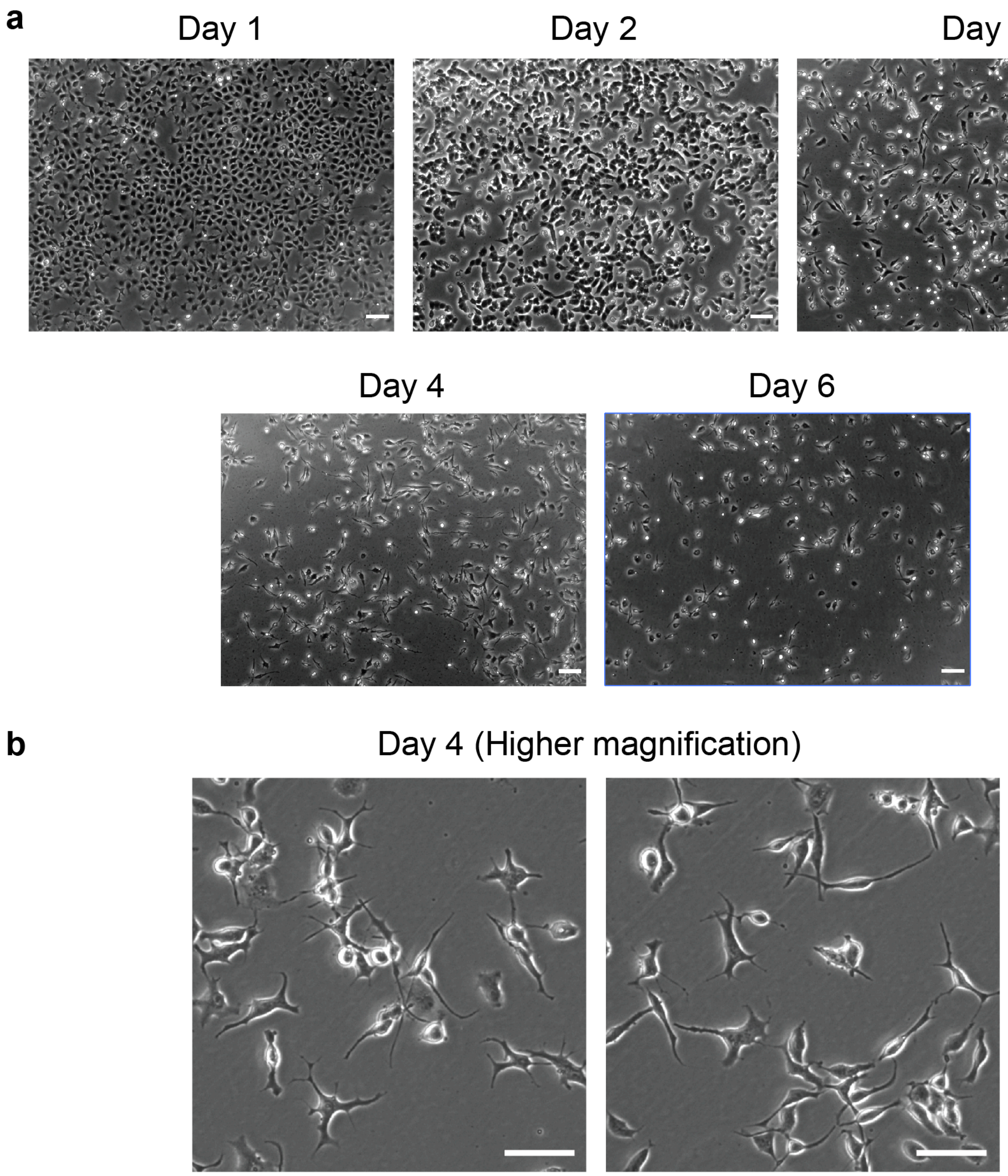


Supplementary Figure S8. Morphology of MG6.4 over different days of Dox induction. (**a**) Images showing morphologies of MG6.4 from day 1 to day 6. Scale bar: 100 μm. (**b**) MG6.4 on Day 4 imaged with higher magnification (Scale bar: 100 μm).

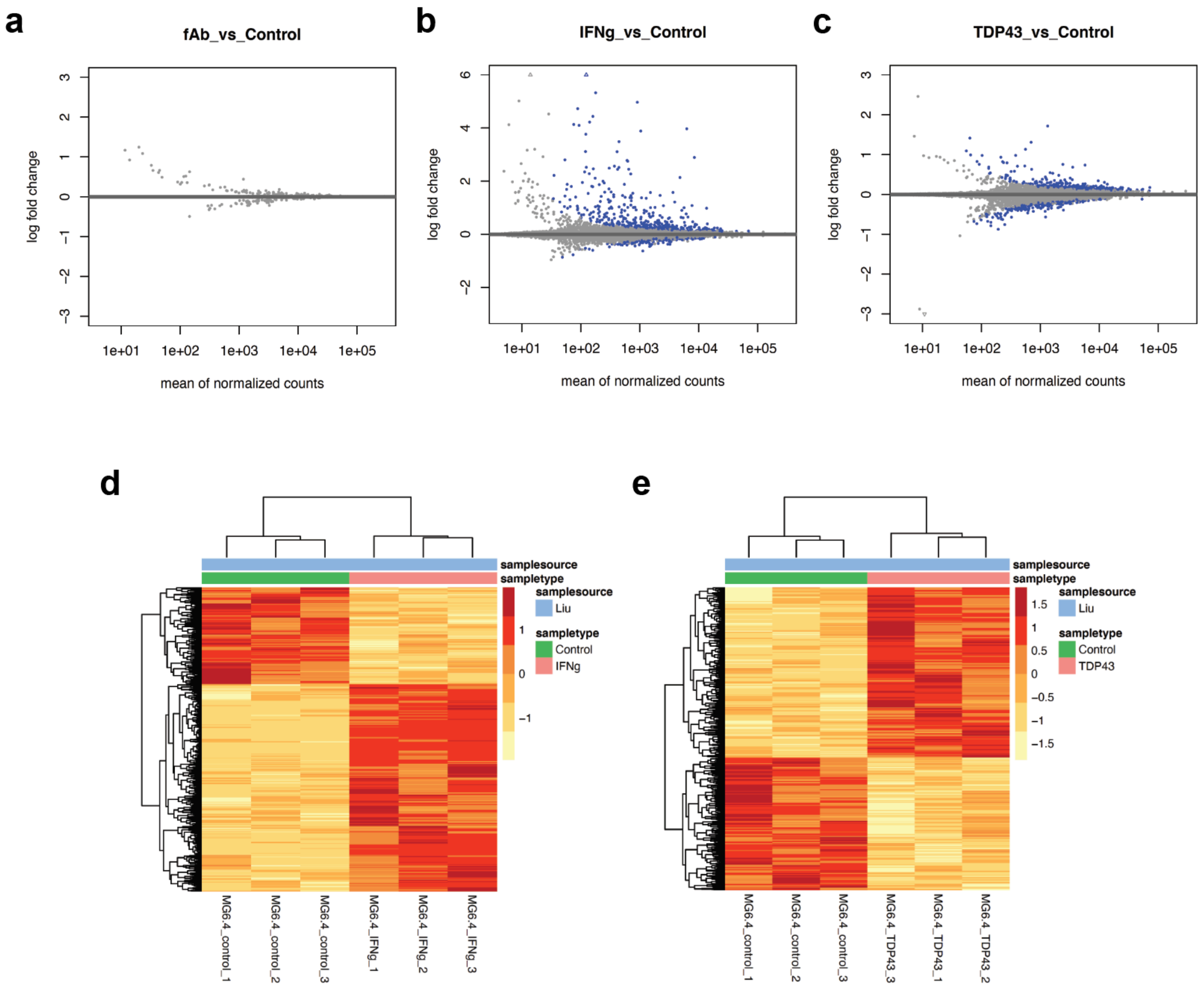


Supplementary Figure S9. (**a-c**) MA plot from DESeq2 analysis of fAβ, IFNγ or TDP-43 treated MG6.4. (**d-e**) Heatmap showing all differentially expressed genes (adjusted p-value < 0.05) between control or stimulated groups.

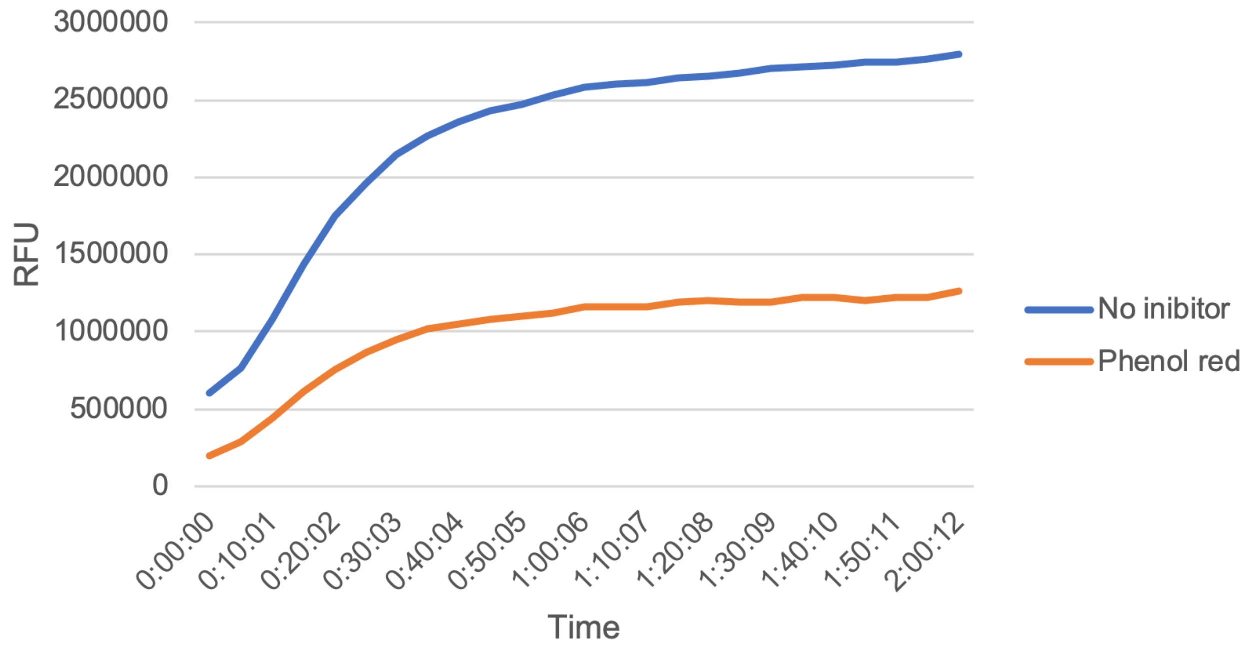


Supplementary Figure S10. Aβ fibrillation experiments with SensoLyte Thioflavin T β-Amyloid (1-42) Aggregation Kit. Phenol red-treated group was included as a known inhibitor control according to manufacturer’s instruction.

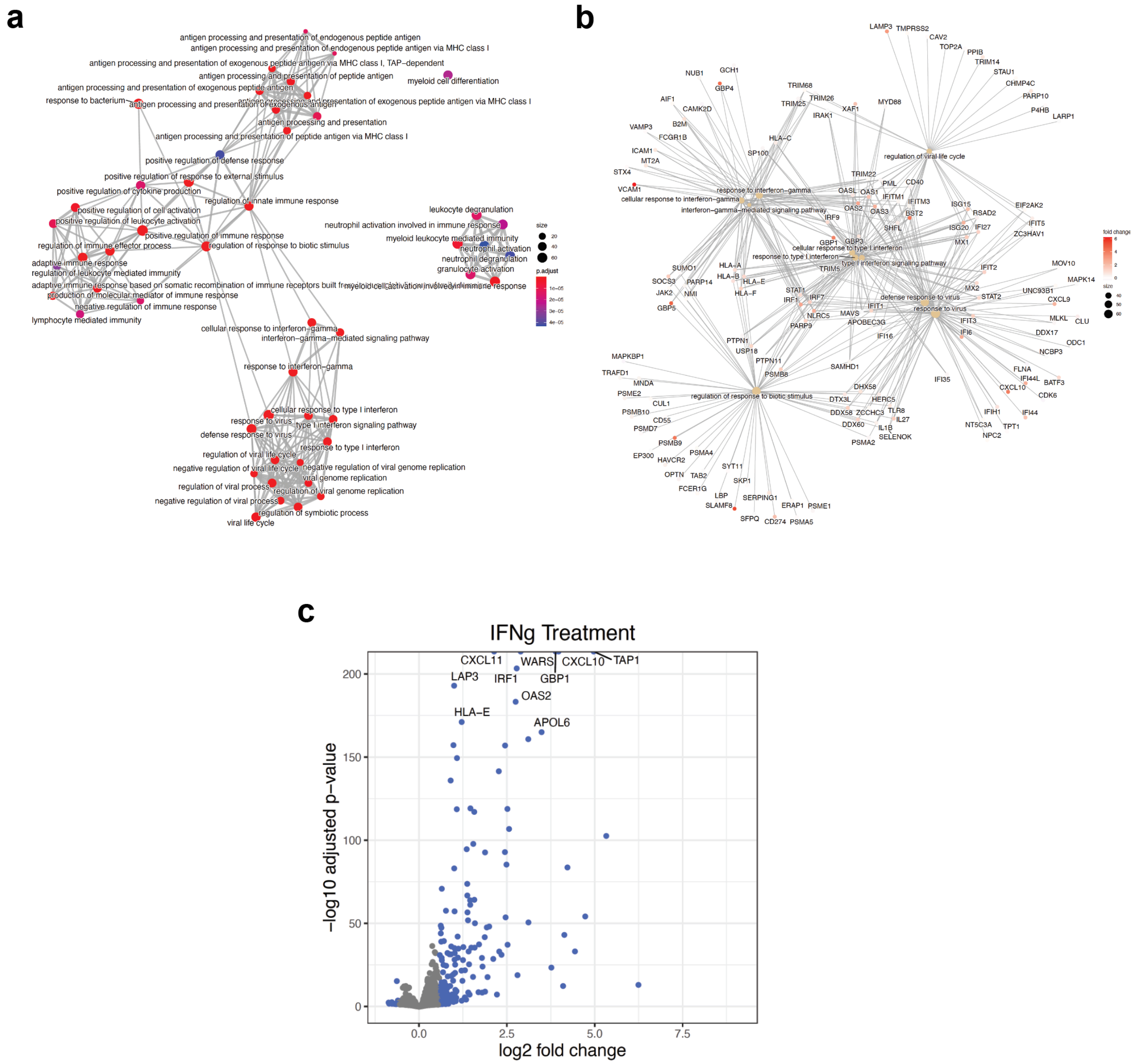


Supplementary Figure S11. Analysis of differentially expressed genes induced by IFNγ in MG6.4. (**a**) Functional module enrichment map showing overlapping gene sets. (**b**) Category netplot showing fold-change of genes associated top significant gene modules. (**c**) Volcano plot showing top differentially expressed genes.

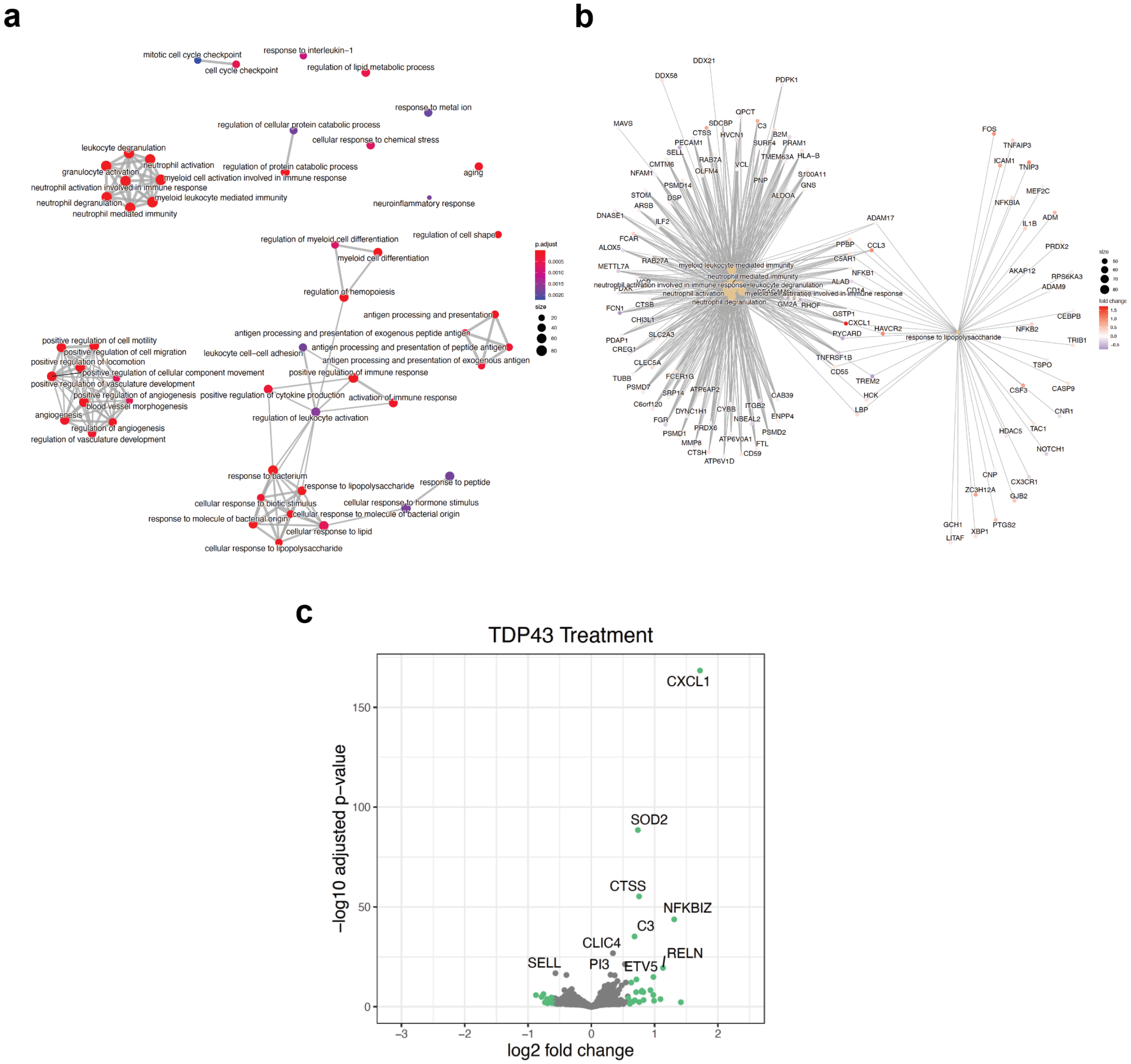


Supplementary Figure S12. Analysis of differentially expressed genes induced by TDP43 in MG6.4. Panel annotation same as Supplementary Figure S11.

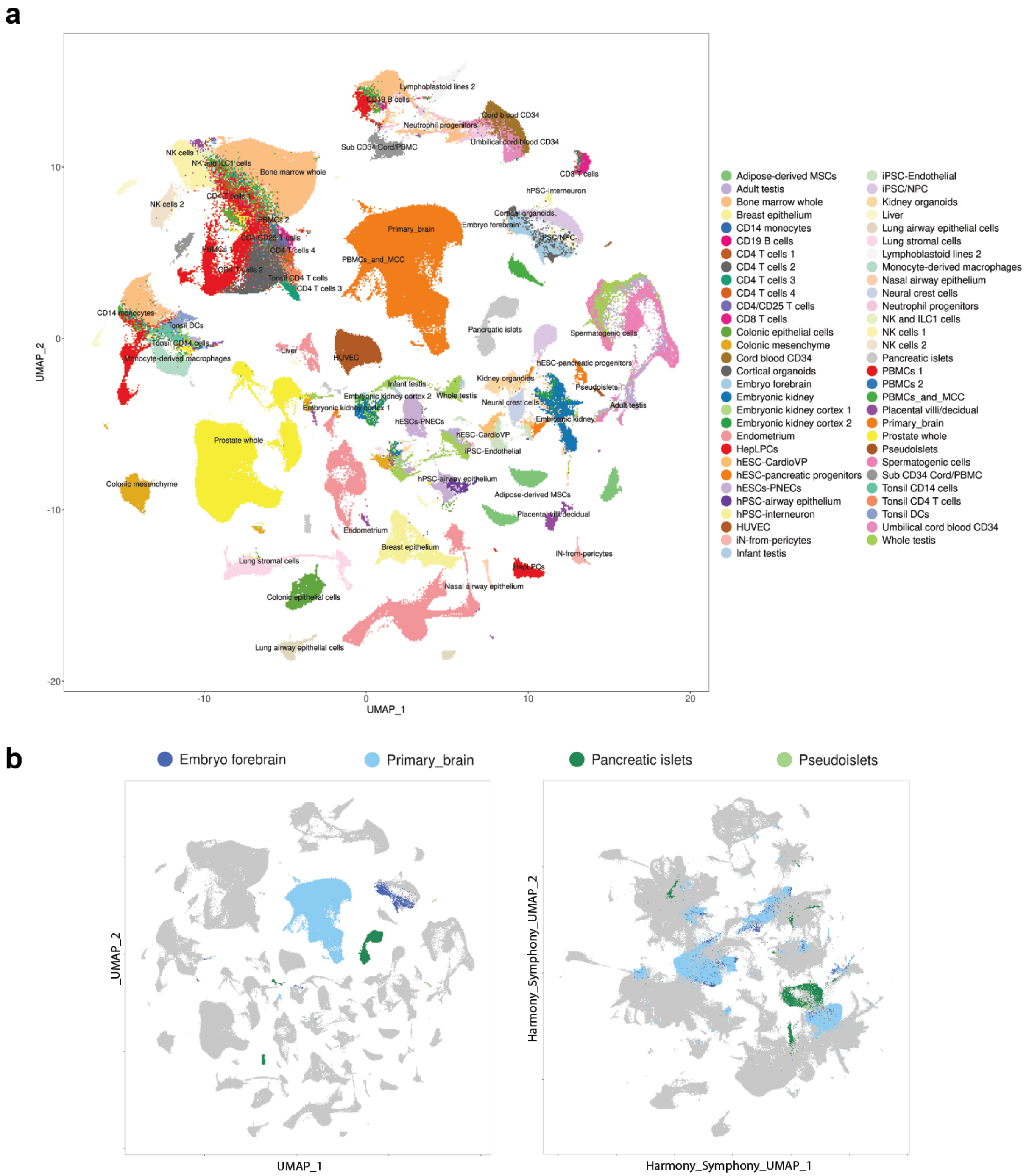


Supplementary Figure S13. Single-cell atlas creation and integration. (a) UMAP visualization of un-integrated single-cell atlas. (b) Demonstration of integration by co-localization of “Primary brain” with “Embryo forebrain”, and “Pancreatic islets” with “Pseudoislets” datasets.

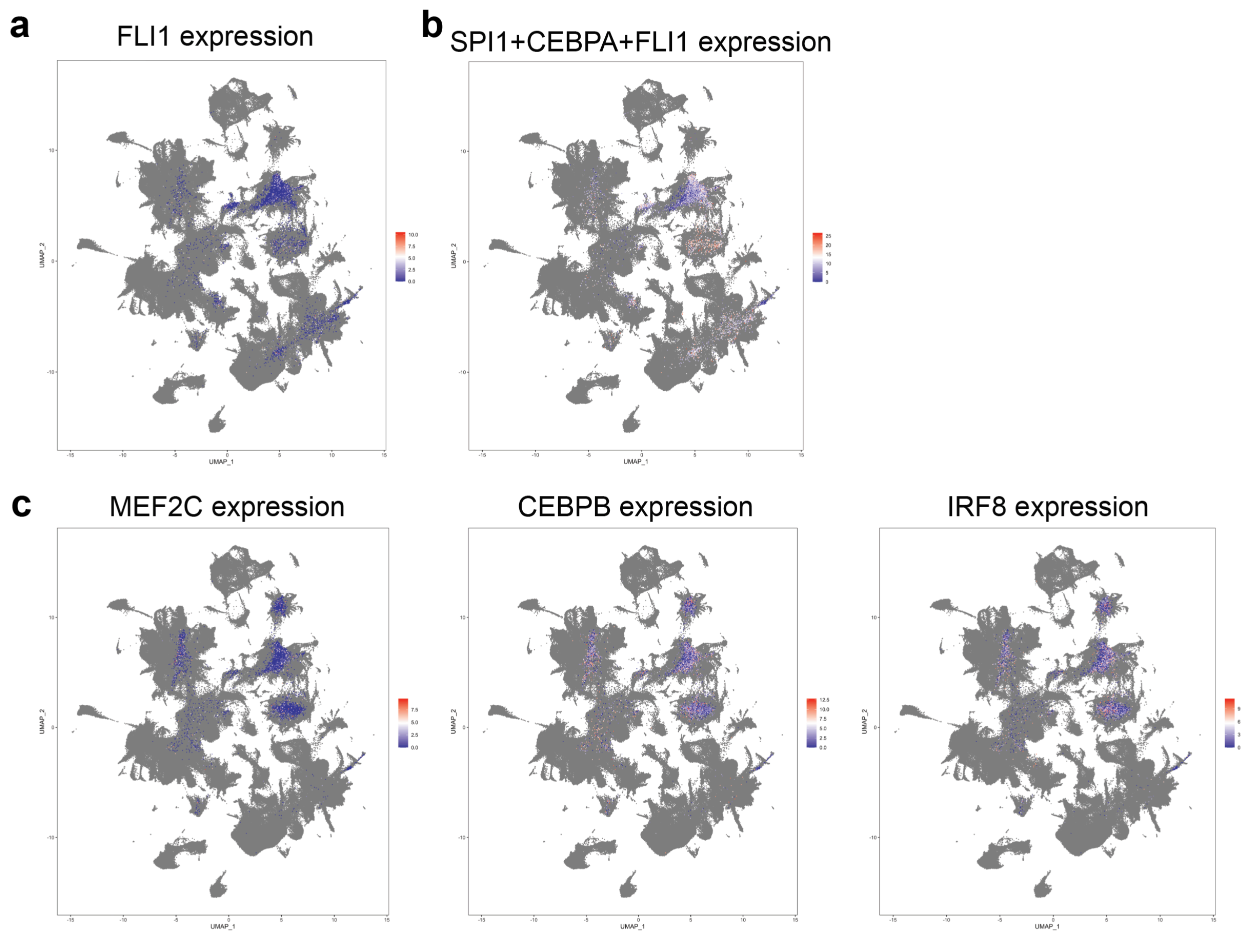


Supplementary Figure S14. Expression of additional TF barcodes in the two screens plotted on single-cell atlas mapping. (a) FLI1 barcode expression in the first pooled screen. (b) Combined expression of SPI1, CEBPA, and FLI1 barcodes. (c) MEF2C, CEBPB, and IRF8 barcode expression in the second pooled screen.

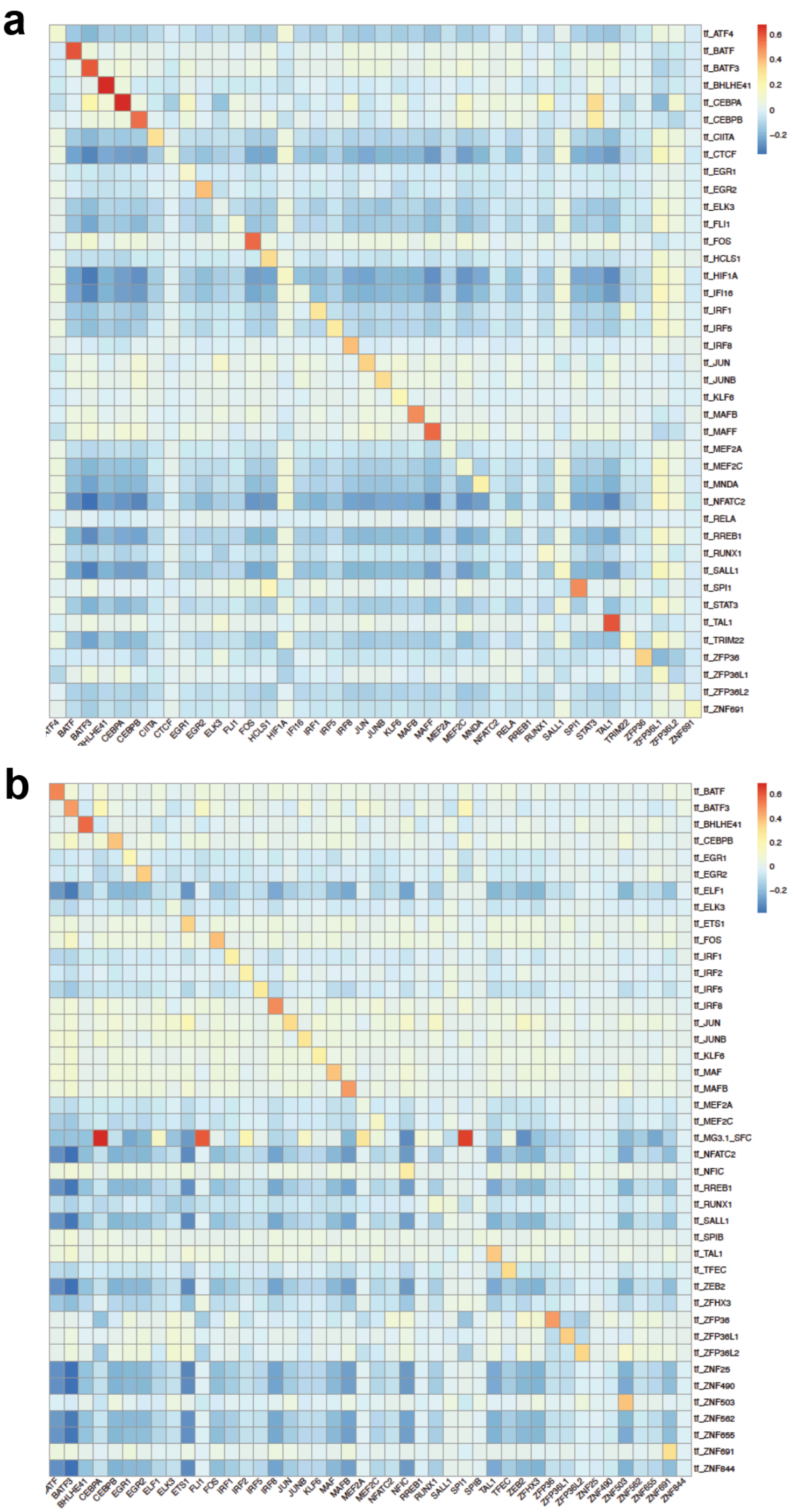


Supplementary Figure S15. Spearman correlation between barcode amplicon sequencing counts and scRNA-seq counts for all the TFs in in (**a**) the first or (**b**) the second pooled screen.

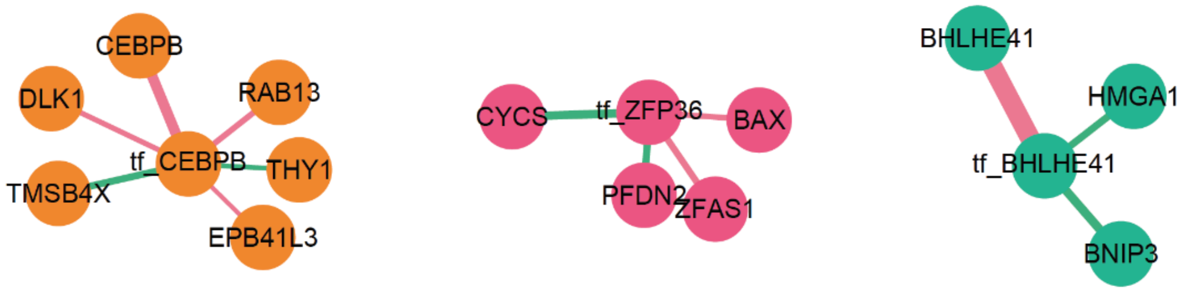


Supplementary Figure S16. Additional sub-networks from the first pooled screen.

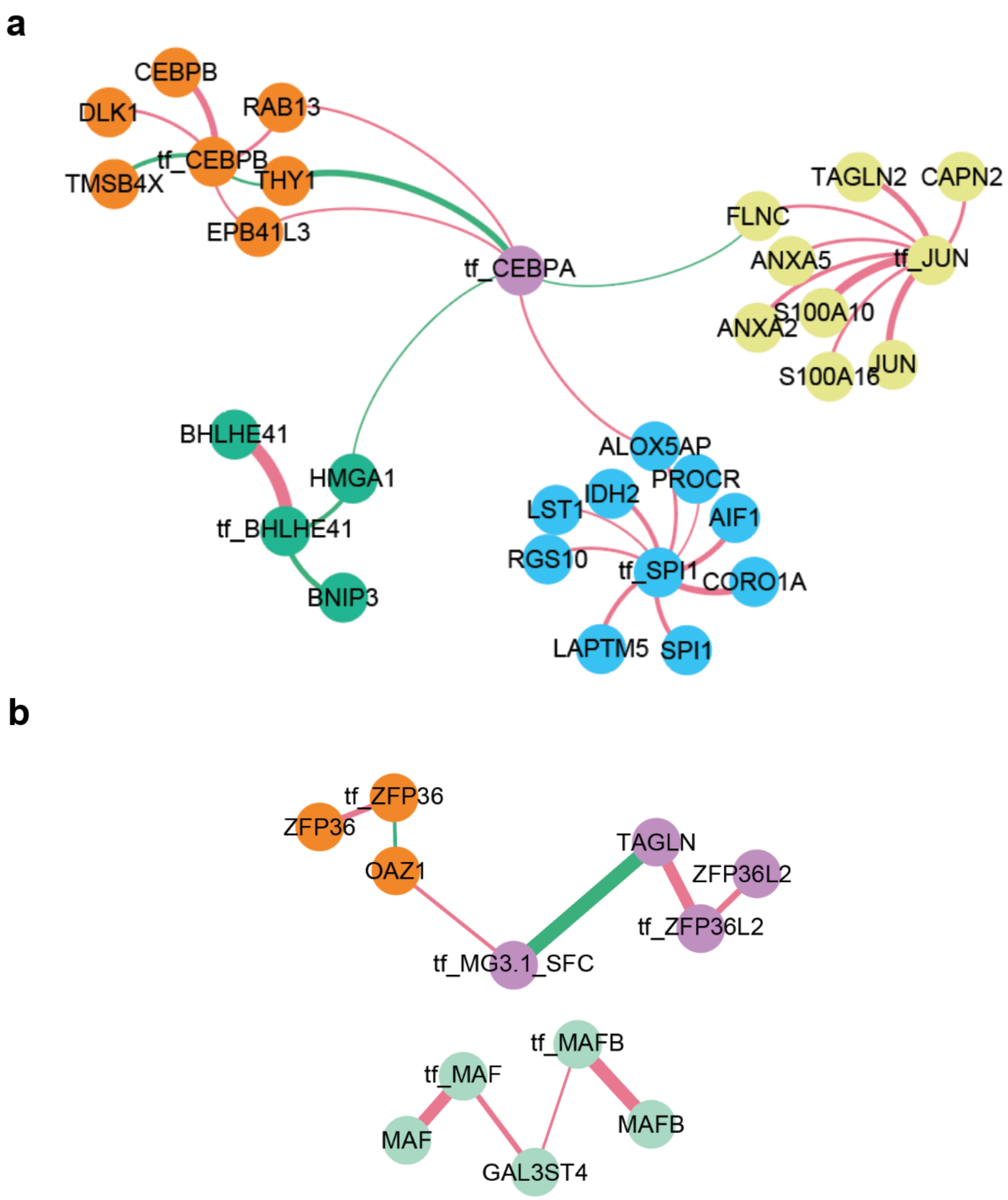


Supplementary Figure S17. Genes that had edges connected to more than one TFs in (**a**) the first or (**b**) the second pooled screen.
